# Towards Convergence in Folding Simulations of RNA Tetraloops: Comparison of Enhanced Sampling Techniques and Effects of Force Field Corrections

**DOI:** 10.1101/2021.11.30.470631

**Authors:** Vojtěch Mlýnský, Michal Janeček, Petra Kührová, Thorben Fröhlking, Michal Otyepka, Giovanni Bussi, Pavel Banáš, Jiří Šponer

## Abstract

Atomistic molecular dynamics (MD) simulations represent established technique for investigation of RNA structural dynamics. Despite continuous development, contemporary RNA simulations still suffer from suboptimal accuracy of empirical potentials (force fields, *ffs*) and sampling limitations. Development of efficient enhanced sampling techniques is important for two reasons. First, they allow to overcome the sampling limitations and, second, they can be used to quantify *ff* imbalances provided they reach a sufficient convergence. Here, we study two RNA tetraloops (TLs), namely the GAGA and UUCG motifs. We perform extensive folding simulations and calculate folding free energies (ΔG_fold_) with the aim to compare different enhanced sampling techniques and to test several modifications of the nonbonded terms extending the AMBER OL3 RNA *ff*. We demonstrate that replica exchange solute tempering (REST2) simulations with 12-16 replicas do not show any sign of convergence even when extended to time scale of 120 μs per replica. However, combination of REST2 with well-tempered metadynamics (ST-MetaD) achieves good convergence on a time-scale of 5-10 μs per replica, improving the sampling efficiency by at least two orders of magnitude. Effects of *ff* modifications on ΔG_fold_ energies were initially explored by the reweighting approach and then validated by new simulations. We tested several manually-prepared variants of gHBfix potential which improve stability of the native state of both TLs by up to ~2 kcal/mol. This is sufficient to conveniently stabilize the folded GAGA TL while the UUCG TL still remains under-stabilized. Appropriate adjustment of van der Waals parameters for C-H…O5’ base-phosphate interaction are also shown to be capable of further stabilizing the native states of both TLs by ~0.6 kcal/mol.

## INTRODUCTION

RNA molecules are present in most viruses and all other living organisms and form dynamics ensembles of many interchanging structures, which play critical roles in a wide range in biological processes.^1–7^ The astonishing multitude of conformations makes RNA structural prediction a difficult task. At present, the static structure of the RNA molecule can be determined/predicted by a number of different experimental and theoretical methods, with variable accuracy.^8^ However, RNA structures are not static and various aspects of RNA structural dynamics, including rarely populated but biochemically relevant conformations, are often important for the function. Atomistic description of structural dynamics of RNA is a major challenge.^8^ Experimental techniques are severely limited in characterization of dynamic motions and thus, computational methods seem to be the ideal complementary technique to study the structure and dynamics of RNA molecules. Valuable insights into RNA structural dynamics can be obtained by atomistic molecular dynamics (MD) simulations.^8^

Standard MD simulations are useful for exploring RNA flexibility, but are limited by the accessible timescale, which now typically reaches lengths of 1-100 microseconds.^8–12^ This timescale should be sufficient to rather reasonably sample dynamics within the main basins on the folding landscape of RNA molecules but does not allow to sample larger structural rearrangements. This obstacle can be overcome by enhanced sampling methods, which allow simulated timescales to be effectively extended in order to probe biologically-relevant conformational changes and chemical reactions.^13^ These methods use several principles to enhance sampling, which differ in the amount of prior information about the energy landscape that is required. While enhanced sampling methods work well for the simplest model systems their efficacy for more complex molecules is often a matter of debate and may vary in a system-method-specific manner.^8, 13–18^

Another fundamental factor determining the usability of the MD technique to study RNA structural dynamics is the quality of the employed empirical potentials (*force fields, ffs*). MD simulations are now routinely used to study dynamics, folding and ligand binding, but the correct description of structure and dynamics of RNA molecules using *ff*s is more problematic than in the case of proteins.^8^ Sampling problems alongside limitations arising from empirical potentials are well-known in the community but their impact on the MD simulation studies is often significantly underestimated.^8, 19, 20^

Compact folded RNA molecules are typically well-described by modern *ff*s on a microsecond timescale when starting simulations from established high-resolution experimental (native) structures and not departing from the basin of the native state. However, the stability of folded RNAs on longer timescales is affected by the free-energy balance between folded, misfolded and unfolded states. Therefore, a common way to probe the accuracy of *ff*s and to identify their major problems (and generally validate them) is to use enhanced-sampling techniques. Considering RNA simulations, replica exchange (RE) methods and (well-tempered) metadynamics (MetaD) are among the most popular enhanced-sampling techniques. They have been extensively applied in folding studies of small RNA tetraloops (TLs), i.e., short RNA stem-loop hairpins with four unpaired nucleotides closed by canonical A-RNA double helix (Figure 1).^21–32^ TLs play a key role in the control of RNA folding, tertiary structure formation, and interactions with other biomolecules.^33–39^ They are ideal motifs to study structural dynamics and folding due to their small size, thermodynamics, and the large amount of experimental data from both Nucleic Magnetic Resonance (NMR) and X-ray crystallography.^40–44^ Many TLs possess a clear dominant folded topology (native state) that is well characterized by a set of signature molecular interactions determining their consensus sequence.^45^ UNCG and GNRA (N and R stands for any and purine nucleotide, respectively) are the most abundant such TLs families. They are shaped-up by intricate balance of diverse molecular interactions and backbone substates, placing high demands on the parametrization of *ff*s.^8, 32, 46^ The 8-mers (stem with two base pairs) are ideal targets for computational modeling as those structures in solution are in dynamic temperature-dependent equilibrium between folded and unfolded conformations. Thus, the *ff*s should capture not only the native state, but also its balance with the unfolded ensemble.^40–43^

**Figure 1:**
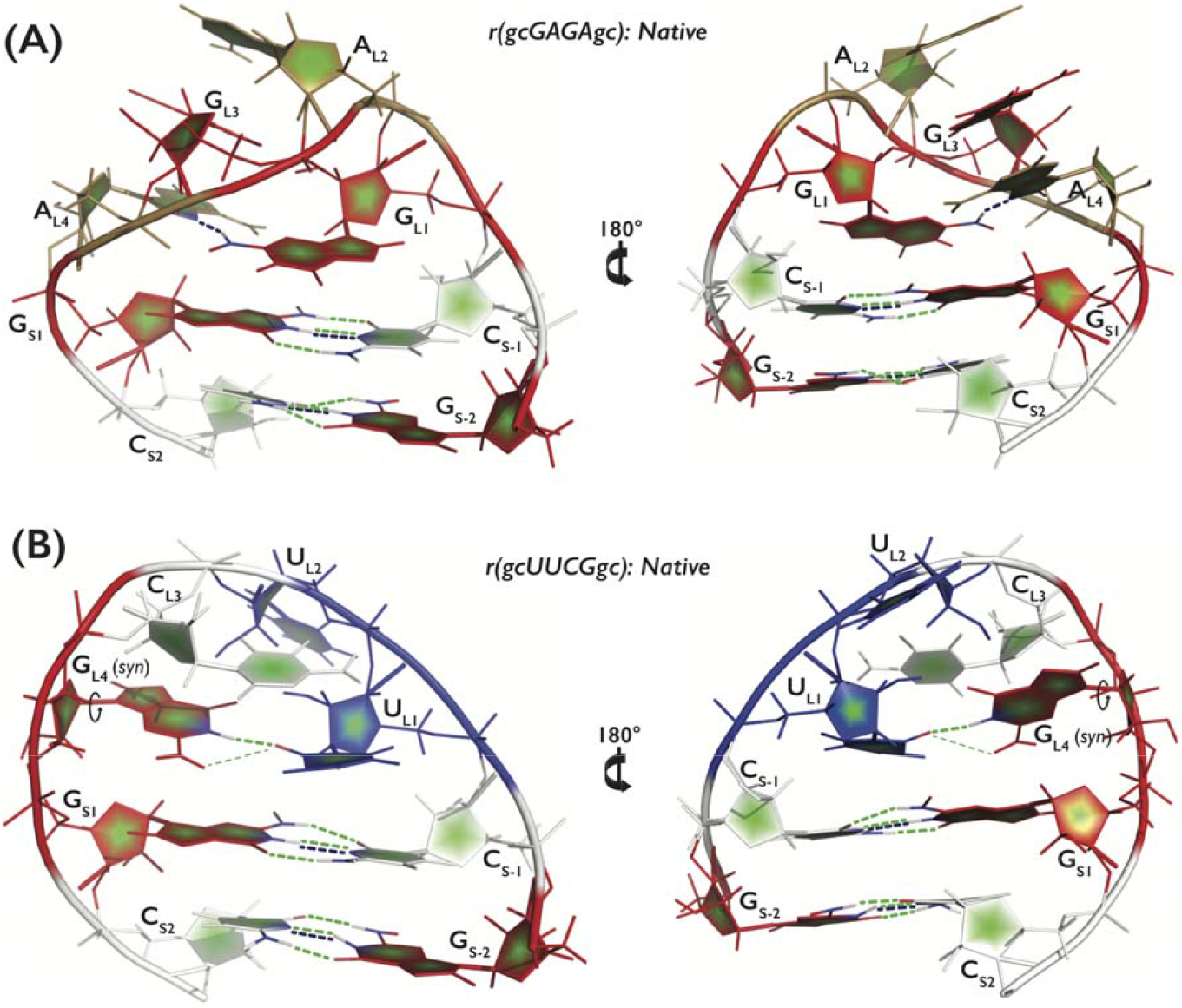
Tertiary structures and detail overview of the r(gcGAGAgc) and r(gcUUCGgc) TLs in their native conformations (see analogous Figure S1 in Supporting Information highlighting signature interactions for both TLs). A, C, G and U nucleotides are colored in sand, white, red, and blue, respectively. The figure compares stabilizing effect of gHBfix_1.0_ (blue dashed lines) and gHBfix_0.5-0.5_ (green dashed lines) potentials on the native state of both TLs. (A) GAGA TL is stabilized by ~3 kcal/mol by both gHBfix_1.0_ (three H-bonds) and gHBfix_0.5-0.5_ (six H-bonds). (B) UUCG TL is stabilized by ~3.5 kcal/mol by the gHBfix_0.5-0.5_ potential (seven or even eight H-bonds due to the bifurcated G_L4_(N1H/N2H)…U_L1_(O2) interaction shown by the thin green dashed line and observed during MD simulations32) and only by ~2 kcal/mol (two H-bonds) by the gHBfix_1.0_ potential. Because the gHBfix potential is as general as all the other *ff* terms it affects all regions of the free energy surface where the specified types of interactions occur.^30^

In the past, various variants of the seminal Cornell et al. (AMBER) parametrization^47^ have been dominantly used to simulate RNA molecules. The most common way to tune performance of the *ffs* are modifications of the dihedral potentials.^29, 48–56^ However, it is increasingly apparent that tuning of dihedral potential has limitations^8^ and more attention has been paid to modifications of the non-bonded *ff* terms. The approaches include basic simple modification of the vdW parameters, reparametrized atomic charges of RNA residues and development of parameters for explicit water models and counter ions.^22, 27, 28, 57, 58^ All the above modifications rely on changes of the parameters of the existing *ff* form. While some promising results have been suggested, subsequent studies indicated that partial improvements for some systems may cause deteriorations for other systems, i.e., side-effects.^8, 30, 59^ This indicates that the present *ff* form is close to the limits of its tunability.

Thus, an alternative possibility is to target just selected (specific) interactions, and in this way to increase the flexibility of parametrization. Popular way to improve *ffs* is to change the pairwise vdW parameters by usage of the so-called nonbonded fix (NBfix) term,^60^ which overrides the universal vdW combination rule for specific pairs of interacting atoms.^22, 27^ Ultimately, one can add new simple terms that are uncoupled from the existing *ff* terms. As such, we recently introduced an additional H-bond potential, named general H-bond fix potential (gHBfix),^24, 30^ which was designed for versatile and well-controllable fine-tuning of specific pairwise H-bond interactions.

The first set of gHBfix parameters led to a substantial improvement of GAGA TL folding.^30^ However, the description of the UUCG TL remains a challenge. Generally, the native state of UUCG TL is lost during long MD simulations and folding simulations mostly indicated a large free-energy disbalance between native/folded and misfolded/unfolded states. ^21–25, 29, 30, 32, 61, 62^ Recent large-scale quantum-chemical (QM) analysis and extensive simulations suggested that the imbalance in the description of the UUCG TLs is due to a concerted effect of multiple *ff* inaccuracies that are coupled and amplifying each other.^32^ The transitory part of energy landscape bridging the well-defined native UNCG conformation with the remaining parts of the landscape appears to be very narrow, which easily leads to conflicts among different *ff* terms. We also suspected that while the advanced replica exchange solute tempering (REST2)^63^ enhanced sampling protocol that we have used in the parametrization of the gHBfix potential worked very well for single strand RNA tetranucleotides, it did not reach a sufficient convergence in folding simulations of the RNA TLs.^30^ RE methods (and generally all methods based on annealing),^8^ are considered as very reliable due to minimal or no introduced prior information. However, there have also been many works suggesting that efficiency of these protocols may be highly-system dependent and in extreme cases can lead to no sampling enhancement.^14, 17–20, 64–68^

In this work, we address both sampling and *ff* issues simultaneously. We used massive enhanced sampling methods in order to study the folding of r(gcGAGAgc) and r(gcUUCGgc) 8-mer TL sequences. We show that for TLs combining REST2 approach with MetaD (i.e., solute tempering with metadynamics, ST-MetaD) outperforms the previously used REST2 method alone. In fact, the REST2 method, at least with ~16 replicas, does not show any sign of convergence for the TLs, in contrast to tetranucleotides. When simulating the UUCG TL on the 16×120 μs REST2 scale, we have observed only four folding events in the whole replica space. In contrast, the ST-MetaD appears to converge the free energies (ΔG_fold_) on the 5 μs per replica scale (with dozens of folding and unfolding transitions). In addition, the ST-MetaD method can be efficiently combined with reweighting. Simultaneously, we tested effects of various *ff* modifications on ΔG_fold_ energies either indirectly by the reweighting approach or directly via new simulations. The stability of native state of both TLs can be quite well tuned by the external gHBfix potential, which modifies populations of H-bond interactions. We tested a few intuitively-selected combinations of the gHBfix parameters while their systematic optimization will be attempted separately. Smaller, but still significant effect comes from NBFix adjustment of van der Waals parameters for 0BPh (base-phosphate type 0)^69^ C-H…O5’ interaction of both purine and pyrimidine nucleotides. Although our *ff* modifications are still not sufficient to get fully satisfactory description of the UUCG 8-mer folding landscape, so that additional modifications will be needed in future, the results are considerably better converged than in the preceding studies. The native structure of the GAGA TL is shown to be stable with all the tested variants. Thus, we show how the RNA *ff* can be tuned by combination of the added gHBfix potential and small NBfix corrections. Effects of those *ff* modifications can be efficiently probed by appropriate enhanced sampling methods, now with the ST-metaD at quantitative level.

## METHODS

### Starting structures and simulation setup

The initial coordinates of r(gcGAGAgc) and r(gcUUCGgc) 8-mers in unfolded single-stranded states were created as described before.^30^ The topology and coordinates were prepared using the tLEaP module of AMBER16 program package.^70^ Single strands were solvated using a rectangular box of OPC^71^ water molecules with a minimum distance between box walls and solute of 12 Å, yielding ~8000 water molecules added and ~65×65×65 Å^3^ box size for both TLs.

We used the standard AMBER OL3 (known also as χ_OL3_)^47, 48, 50, 72^ RNA *ff* The basic OL3 version was applied with the vdW modification of phosphate oxygens developed by Steinbrecher et al.,^73^ where the affected dihedrals were adjusted as described elsewhere.^51^ AMBER library file of this *ff* version can be found in Supporting Information of Ref. ^24^. This *ff* version is abbreviated as χ_OL3CP_ henceforth. AMBER topologies and coordinates were then converted into GROMACS inputs via PARMED. Simulations were run at ~1 M KCl salt excess using the Joung-Cheatham ion parameters^75^ (K^+^: r =1.705 Å, ε = 0.1937 kcal/mol, Cl^−^: r = 2.513 Å, ε= 0.0356 kcal/mol) at T = 298 K with the hydrogen mass repartitioning^76^ allowing an 4-fs integration time step (see Supporting Information of Ref. ^30^ for other details about the simulation protocol).

### Additional *ff* modifications by gHBfix and NBfix potentials

We used the gHBfix potential,^30^ which is an additional *ff* term developed recently for tuning of H-bonding interactions in nucleic acids; for full description see Ref. ^30^. We mainly tested two different versions of the gHBfix potential. Both versions strengthened base – base H-bonds and weakened sugar – phosphate interactions as suggested by the original paper^30^ but differ in the way how they stabilize base – base interactions. The first version labelled here as gHBfix_1-0_ is identical to the originally suggested version,^30^ which is nowadays marked in the literature as gHBfix19.^32, 77^ It strengthens all –NH…N– interactions by 1.0 kcal/mol, while the –NH…O– interactions are unaffected. The second gHBfix version (gHBfix_0.5-0.5_) strengthened both –NH.N– and –NH…O– interactions by 0.5 kcal/mol. Figure 1 depicts, which H-bonds are stabilized by gHBfix_1.0_ and gHBfix_0.5-0.5_ in the native state of both TLs. Additionally, we tested a more complex version of the gHBfix potential developed for the UUCG TL. The gHBfix_UNCG19_ version does not only modify base donor – base acceptor and sugar-phosphate H-bonds but in addition sugar donor – base acceptor H-bonds are strengthened and base donor – sugar acceptor and sugar – sugar H-bonds weakened (see Ref. ^32^ for full description).

We also performed simulations with *ff* corrections using the so-called nonbonded fix (NBfix) approach^60^ modifying the pairwise vdW parameters via breakage of combination (mixing) rules. Via NBfix we reduced the minimum-energy distance of Lennard-Jones potential (i.e., *R_ij_* parameter) for the –H8…O5’– and –H6…O5’– pairs, i.e., between H5 – OR and H4 – OR atom types; by 0.25Å to 2.8808 Å for purine (NBfix_0BPh-_pur__) and 2.9308 Å for pyrimidine (NBfιx_0BPh-_pyr__) nucleotides, respectively. The OR atom type includes both O5’ and O3’ bridging phosphate oxygens but the involvement of the O3’ atom in the 0BPh interactions is marginal. Depths of potential wells (*ε_ij_* parameters) were kept at their default value of 0.0505 kcal/mol. In other words, we decreased the repulsion between H8 and H6 atoms of all purine and pyrimidine bases and O5’ oxygens of phosphates. While we prefer to tune H-bonding interactions via the gHBfix *ff* term, NBfix is more straightforward to correct excessive short-range atom-atom repulsions.^22, 27, 60^

For the sake of completeness, in one REST2 simulation of UUCG TL with the gHBfix_UNCG19_ potential, we also generally reduced vdW radii of all non-polar H atoms, i.e., vdW radii of H1, H4, H5 and HA atoms were universally changed to 1.2 Å. Reduction of non-polar H atoms was applied together with the NBfix_0Bph-_pur__ correction as we followed the setup from our earlier works.^30, 32^ We are, however, not widely using this modification since its effect is unclear; we rather assume that this modification has negligible effect on the folding landscape.

### Enhanced sampling

Well-tempered MetaD^78, 79^ in combination with REST2 method^63^ was used to accelerate sampling and transitions between native and other (unfolded and misfolded) states. The method is abbreviated here as ST-MetaD and was firstly applied by Camilloni et al. for G-helix protein folding.^80^ While MetaD helps to overcome large energy barriers, REST2 improves the sampling and allows to reduce dramatically the number of replicas in comparison with parallel-tempering (T-REMD) MetaD approach.^25, 81, 82^ For both TLs, 12 replicas starting from unfolded single strands were simulated in the effective REST2 temperature range of 298–497 K (the scaling factor λ values ranged from 1.0 to 0.59984) for 5 μs per replica while some simulations were prolonged to 10 μs (Table S1 in the Supporting Information). The average acceptance rate was ~30% for both TLs.

The *ε*RMSD metric^83^ was used as a biased collective variable.^25^ We used *ε*RMSD with an augmented cutoff (set at 3.2) for biasing, which was shown to allow forces to drive the system towards and away from the native state even when nucleobases are far from each other.^25^ The ĉRMSD with the standard cutoff (set at 2.4)^83^ was used for monitoring states throughout simulations and following population analysis required for calculation of free energies (see below). Reference native structures for TLs were taken from our previous work^30^ and were determined based on simultaneous observation of both a low *ε*RMSD from the native state and the presence of all native H-bonds, i.e., the base pairing in the stem (two GC base pairs) and signature interactions of the loop^45^ (Figure S1 in Supporting Information). Presence of a H-bond was inferred from a hydrogen-acceptor distance within cutoff 2.5 Å. ST-MetaD simulations were carried out using GPU-capable version of Gromacs2018^84^ in combination with PLUMED 2.5.^85, 86^ Sampling of each replica was enhanced by Gaussians deposited every 1 ps (i.e., 250 steps) with a height of 0.118 kcal/mol, and Gaussian width (*σ* parameter) set to 0.1. ST-MetaD rescaled the height of the Gaussians with a bias factor (T +ΔT)/T of 15.

We also performed one extra simulation with the “standard” REST2 protocol^63^ for each TL. Those were run at 298 K (the reference replica) with 16 replicas. While the r(gcGAGAgc) simulation was run for 13 μs per replica, the r(gcUUCGgc) simulation was prolonged to enormous 120 μs per replica. The λ values ranged from 1.0454 to 0.59984 and those values were chosen to maintain an exchange rate above 20%. The effective solute temperature ranged from 285 K to ~500 K. Details about REST2 settings can be found elsewhere.^30^ See Table S1 in Supporting Information for list of all performed simulations.

### Estimated folding free energy balances of 8-mer TLs

The εRMSD threshold (calculated with standard cutoff) separating the folded and un(mis)folded states was set at value of 0.7 for both TLs.^25^ We calculated folding free energy difference (ΔG_fold_) from populations of the native structure and other conformations. Free energy differences were obtained as: *ΔG_fold_ = –RT[(ln(Σ_i,folded_ W_i_) – In[(ln(Σ_i,un(mis)foided_ W_i_)]*, where *W_i_*, stands for weighting factor of i-th snapshot considering final time-averaged bias potential from each replica and weight of the particular snapshot within the replica space with respect to the reference replica ensemble. The free energy calculation thus uses information from all replicas albeit only few lowest replicas are able to contribute significantly due to weighting factors. Time-averaging of bias potentials is routinely used for non-well-tempered MetaD^87^ and is, in principle, not necessary for well-tempered MetaD, where the updated rate of the bias potential decreases as 1/t, where t is the simulation time. However, it was recently shown that time-averaging can significantly speed up convergence of estimated ΔG_fold_ energies as instantaneous estimates from final bias potentials were significantly affected by fluctuations.^88^ Figures 2 and 3 show that instantaneous estimates of ΔG_fold_ energies indeed fluctuate, whereas those calculated from time-averaged bias potentials are more stable.

**Figure 2:**
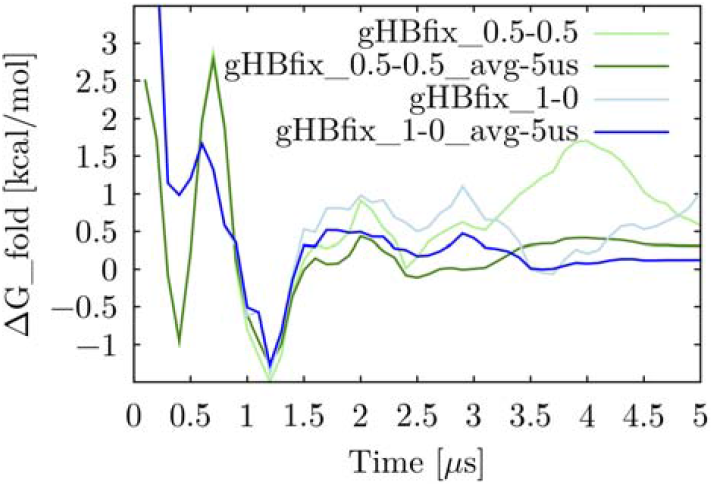
Fluctuations of ΔG_fold_ energies from 5 μs-long ST-MetaD simulations of r(gcGAGAgc) TL. Behavior with gHBfix_0.5-0.5_ (green curves) and gHBfix_1-0_ (blue curves) potentials are shown. Energies were calculated by using bias potentials over 100 ns-long windows from the reference replica only (the lowest REST2 replica corresponding to 298 K), i.e., 50 points, from the respective simulations. Instantaneous estimates of ΔG_fold_ energies (shaded lines) fluctuate, whereas those calculated by time-averaged bias potentials are converged (bold lines labelled as “avg” in the legend). Note that *ε*RMSD with augmented cut-off (3.2 Å) was used for this specific analysis, where we focused on convergence instead of absolute values of ΔG_fold_ energies. Thus, final ΔG_fold_ energies displayed on the plot are slightly different from the values reported in the text, which were obtained from bias potentials of all twelve replicas and by ĉRMSD with standard cut-off more accurately identifying the native states (see Methods and Table 1).

**Figure 3:**
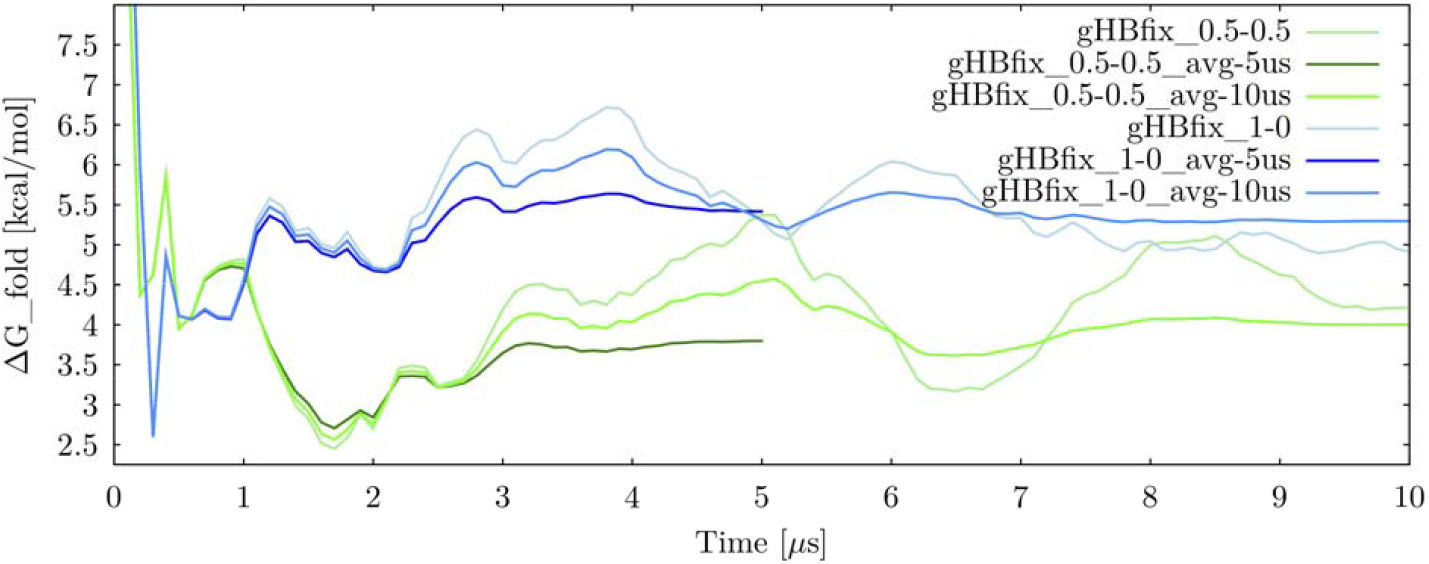
Fluctuations of G_fold_ energies from 10 μs-long ST-MetaD simulations of r(gcUUCGgc) TL. Behavior with gHBfix_0.5-0.5_ (green curves) and gHBfix_1-0_ (blue curves) potentials are shown for comparison. Energies were calculated by using bias potentials over 100 ns-long windows from the reference replica only (see Figure 2 legend for more details). ΔG_fold_ energies calculated by time-averaged bias potentials (bold lines labelled as “avg” in the legend) using either data from 5 μs-long or prolonged 10 μs-long simulations are comparable, showing good convergence of the 5 μs-long timescale.

We also estimated effects of various *ff* modifications during ST-MetaD simulations by the reweighting algorithm. Reweighting allows to calculate the unbiased (or biased) probabilities from the biased (or unbiased) probabilities and, consequently, reconstructing free energy profiles, e.g., ΔG_fold_ balances, from the (un)biased probabilities.^90, 91^ Reweighting was performed using the final averaged bias potential with snapshots only from the reference replica (the lowest REST2 replica corresponding to 298 K).

The convergency of ST-MetaD simulations was assessed by computing populations and ΔG_fold_ energies from final time-averaged bias potentials after 2 μs, 3 μs, 4 μs, and 5 μs-long windows (Figure S2 in Supporting Information). Statistical errors were calculated by estimating higher and lower boundaries of populations of the native structure. Those were obtained from concatenated trajectories and bootstrapping with 16 and 32 blocks (for 5 μs-long and 10 μs-long simulations, respectively), where block size was 312.5 ns. We also tested different size of blocks and statistical errors were always similar and lower than ~0.25 kcal/mol (see Figure S3 in Supporting Information for evolution of estimated errors). Nonetheless, we would like to stress that the bootstrap analysis takes into account only the statistical fluctuations of the folded state population in time domain, while it does not consider the convergence of the final (averaged) bias potential used in the calculation of the snapshot weights. Therefore, true uncertainties of calculated free energy differences might be larger than those estimated by bootstrap analysis.

The convergence of the “standard” REST2 simulations was estimated by sophisticated bootstrapping protocol introduced in Ref. with resampling in both time and replica domains. That enhanced bootstrap procedure enables distinguishing between cases where the native state is formed only in just one or very few replicas versus cases when the native state is repeatedly formed in most or all replicas. If the latter is not true then the very few (sometimes just one) folding events result into huge uncertainty; cf. the large statistical errors in our previous attempts to fold TLs by using the REST2 approach.^30^ Hence, we did not calculate folding ΔG_fold_ energies from populations of the native structure observed in REST2 simulations when the estimated statistical error would be larger than the estimated population of the native state, effectively including the *zero* population into the target confidence bootstrap interval (Table 1). We notice that any quantification of the error could be subject to significant underestimation in cases were the number of transitions between the relevant states is too low or if some portions of the energy landscape are not sampled.

**Table 1:** Calculated folding free energy (ΔG_fold_) and population of native state (p_native_) at 298 K for r(gcGAGAgc) and r(gcUUCGgc) TLs.^a^

| TL | *ff* modification | $\Delta G_{\text{fold}}$<br>(kcal/mol)<sup>b</sup> | $p_{\text{native}}$, 298 K (%) |
| --- | --- | --- | --- |
| r(gcGAGAgc) | gHBfix<sub>0.5-0.5</sub> | $0.29 \pm 0.17$ | $37.97 \pm 6.52$ |
| r(gcGAGAgc) | gHBfix<sub>1-0</sub> | $0.14 \pm 0.16$ | $44.13 \pm 6.65$ |
| r(gcUUCGgc) | gHBfix<sub>0.5-0.5</sub> | $3.90 \pm 0.11$ | $0.14 \pm 0.03$ |
| r(gcUUCGgc) | gHBfix<sub>1-0</sub> | $5.42 \pm 0.10$ | $0.01 \pm 0.00$ |
| r(gcUUCGgc) | gHBfix<sub>0.5-0.5</sub>_NBfix<sub>0BPh-pur</sub> | $4.06 \pm 0.15$ | $0.11 \pm 0.03$ |
| r(gcUUCGgc) | gHBfix<sub>1-0</sub>_NBfix<sub>0BPh-pur</sub> | $5.48 \pm 0.08$ | $0.01 \pm 0.00$ |
| r(gcUUCGgc) | gHBfix<sub>0.5-0.5</sub>_NBfix<sub>0BPh-pyr</sub> | $2.32 \pm 0.11$ | $2.01 \pm 0.35$ |
| r(gcUUCGgc) | gHBfix<sub>1-0</sub>_NBfix<sub>0BPh-pyr</sub> | $4.67 \pm 0.09$ | $0.04 \pm 0.01$ |
| r(gcUUCGgc) | gHBfix<sub>UNCG19</sub>_NBfix<sub>0BPh</sub><sup>c</sup> | $1.87 \pm 0.12$ | $4.17 \pm 0.82$ |
| r(gcGAGAgc)<sup>d</sup> | gHBfix<sub>1-0</sub> | - | $19.20 \pm 17.32$ |
| r(gcUUCGgc)<sup>e</sup> | gHBfix<sub>UNCG19</sub>_NBfix<sub>0BPh-pur</sub> | - | $4.97 \pm 5.65$ |

<sup>a</sup> All simulations were run with the $\chi_{\text{OL3CP}}$ AMBER RNA *ff* using either the ST-MetaD or REST2 (the last two lines) methods.
<sup>b</sup> Free energies were estimated from 5 μs-long simulations using concatenated trajectories from all 12 replicas and time-averaged bias potentials – see Methods.
<sup>c</sup> More complex version of gHBfix potential (gHBfix<sub>UNCG19</sub>)<sup>32</sup> specifically designed for the UUCG TL (see Methods).
<sup>d</sup> 16 x 13 μs REST2 simulation with the gHBfix<sub>1-0</sub> potential. $\Delta G_{\text{fold}}$ energy was not calculated due to very large statistical error – see Methods.
<sup>e</sup> 16 x 120 μs REST2 simulation with the gHBfix<sub>UNCG19</sub> potential. In addition, vdW radii of all non-polar H atoms were reduced (see Methods). $\Delta G_{\text{fold}}$ energy was not calculated due to very large statistical error.

## RESULTS AND DISCUSSION

Previous studies typically reported significant imbalance of estimated folding free energies (ΔG_fold_) for small RNA TLs, with the simulations underestimating stability of the folded state.^24, 25, 27, 30^ The discrepancy between theory and experiment was primarily attributed to inaccuracies of the common AMBER RNA *ffs* but overall time-scale of simulations and convergence of enhanced sampling methods were also questioned. Here, we provide series of extensive enhanced sampling ST-MetaD simulations of r(gcGAGAgc) and r(gcUUCGgc) RNA TLs, which seem to resolve the issue of convergence and provide some hints on how to improve the *ff*. We calculated the ΔG_fold_ energies and investigated effects of several *ff* modifications on stability of the native state. In particular, we tested effects of two variants of the gHBfix potential and some modifications of pairwise vdW parameters. Those effects were probed either directly in simulations or indirectly by the reweighting approach. In total, we carried out nine ST-MetaD simulations with 12 replicas with a cumulative time of 780 μs and two REST2 simulation with 16 replicas and cumulative time of 2128 μs, so that the study reports more than 2.9 ms of new simulation data (Table S1 in Supporting Information). Additional REST2 simulations for comparison were taken from the preceding works. ^30, 32, 58^ We demonstrate much better convergence of the ST-MetaD approach compared to the sole REST2 method. The results are organized in the following way. We first present the ST-MetaD simulations and analyze the suggested *ff* modifications. Comparison of the ST-MetaD and REST2 protocols is discussed subsequently.

### ST-MetaD simulations of GAGA and UUCG TLs with gHBfix_1-0_ and gHBfix_0.5-0.5_ potentials

Initial ST-MetaD simulations of r(gcGAGAgc) and r(gcUUCGgc) were done with two different versions of the gHBfix potential. We wanted to determine how different stabilization of base – base interactions by the gHBfix affects folding energies. The first version labelled as gHBfix_1-0_ is stabilizing only –NH…N– interactions by 1.0 kcal/mol while –NH…O– interactions are not modified. This version (abbreviated in the literature as gHBfix19^32, 77^) was suggested earlier based on the REST2 simulations of TLs and tetranucleotides. In the second version (labelled here as gHBfix_0.5-0.5_) both –NH…N– and –NH…O– interactions are stabilized by 0.5 kcal/mol (see Methods for details). Thus, each canonical GC base pair (forming three H-bonds) was stabilized by 1.0 kcal/mol and 1.5 kcal/mol by gHBfix_1.0_ and gHBfix_0.5-0.5_, respectively (Figure 1).

The GAGA 8-mer TL simulations predict essentially identical ΔG_fold_ energies of 0.14 ± 0.16 kcal/mol and 0.29 ± 0.17 kcal/mol with gHBfix_1.0_ and gHBfix_0.5-0.5_ potentials, respectively (Table 1). It is not unexpected as both gHBfix_1-0_ and gHBfix_0.5-0.5_ potentials aim to stabilize the native state by 3 kcal/mol (Figure 1). Both calculated ΔG_fold_ energies correspond to the population of native state above 40 % at 298 K, which is in reasonable agreement with experiments.^42, 43^ Importantly, the calculated r(gcGAGAgc) 8-mer ΔG_fold_ values are much closer to the experiment and significantly lower (by ~3.7 kcal/mol) than the one reported previously without the gHBfix (3.9 ± 0.1 kcal/mol).^25,93^ The difference is qualitatively consistent with contributions of gHBfix potentials (i.e., 3.0 kcal/mol and 3.5 kcal/mol by gHBfix_1-0_ and gHBfix_0.5-0.5_, respectively), even though the stem sequence and the water model used in Ref. ^25^ is different from the one simulated here, and the MetaD approach was combined with T-REMD method.^25^

The calculated ΔG_fold_ energies for UUCG TL are in very positive territory (Table 1), which is not surprising considering the notorious *ff* problems in description of this TL. ^21, 25, 27, 30, 32, 54, 57, 94^ However, both calculated ΔG_fold_ energies (3.90 ± 0.11 kcal/mol and 5.42 ± 0.10 kcal/mol with gHBfix_0.5-0.5_ and gHBfix_1-0_ potentials, respectively) are again lower than the ΔG_fold_ energy of 5.5 ± 0.1 kcal/mol reported previously without the gHBfix potential for the r(ccUUCGgg) TL.^25, 93^ The ΔG_fold_ energy with the gHBfix_0.5-0.5_ potential is notably lower (by ~1.5 kcal/mol, Table 1) compared to the gHBfix_1-0_ potential. It is because the gHBfix_0.5-0.5_ potential targets larger stabilization of the UUCG native state than the gHBfix_1-0_ (see Figure 1 for visualization of interactions stabilized by the two gHBfix variants). The two GC base pairs forming a short stem of the 8-mer motif are more stabilized by the gHBfix_0.5-0.5_, which is fully transferred to the actual ST-MetaD calculation. Additionally, the gHBfix_0.5-0.5_ is stabilizing the GU-wobble base pair within the loop (either by single or bifurcated H-bond, Figure 1), while this base pair is not stabilized by the gHBfix_1.0_ potential.

The GAGA TL ΔG_fold_ data did not suggest a difference between gHBfix_0.5-0.5_ and gHBfix_1.0_ potentials because larger stabilization of the stem in gHBfix_0.5-0.5_ with respect to gHBfix_1.0_ potential is partially compensated for by destabilization of the loop. In addition, the stem in GNRA TLs is expected to be more stable than the same stem in UNCG TLs as the 3’-overhang G_L1_ guanine in GNRA TLs should provide larger stacking stabilization than U_L1_ uracil of UNCG TLs.^95^ The gHBfix_1-0_ potential seems to be sufficient for folding of GNRA TLs, while simulations of UNCG TLs with pyrimidine 3’-overhang may, besides revealing instabilities within the loop itself, efficiently unmask possible under-stabilization of short stems in contemporary *ffs*.

In summary, the initial set of calculated ΔG_fold_ energies from ST-MetaD simulations shows that the external gHBfix potential improves their stability compared to the χ_OL3CP_ *ff* alone for both TLs. Both gHBfix versions are sufficient to correctly fold the GAGA TL. On contrary, the calculated ΔG_fold_ energies for UUCG TL are still high and the fold/unfold free energy imbalance is only partially reduced. The gHBfix_0.5-0.5_ version outperforms the original gHBfix_1-0_ (known as gHBfix19) for the UUCG TL.

### UUCG TL folding simulations with NBfix correction of the 0BPh interaction

The NBfix_0BPh-_pur__ correction (see Methods) was tentatively suggested based on QM calculations with the goal to stabilize the canonical UUCG TL sugar – phosphate backbone conformation between the *syn* G_L4_ and neighboring G_S+1_ bases. The correction reduces the –H8(G_S+1_) …O5’(G_S+1_) steric repulsion, which is excessive with the *ff* description and seems to cause flip of the G_S+1_ backbone from the canonical conformation (which is native to the TL in this position) to nonnative *β_g+/γtrans_* G_S+1_ state.^32^ Here, we explicitly tested its effect on the ΔG_fold_ by performing two r(gcUUCGgc) ST-MetaD simulations with gHBfix_1.0_ and gHBfix_0.5-0.5_ potentials coupled with the NBfix_0BPh-_pur__ correction, i.e., the modification applied to all purines.

The NBfix_0BPh-_pur__ correction did not significantly affect the ΔG_fold_ values (Tables 1 and 2). This is somewhat surprising, as preceding standard simulations suggested that the NBfix_0BPh-_pur__ correction alleviates a steric conflict in the folded state. Thus we (i) prolonged all four simulations (both with and without NBfix_0BPh-_pur__ correction) to 10 μs and (ii) carried out cross-reweighting, where the simulations without NBfix were reweighted with the NBfix and simulations with the NBfix were reweighted back for the standard parameters. Calculated ΔG_fold_ energies from the prolonged simulations are 5.29 ± 0.08 kcal/mol, 4.03 ± 0.11 kcal/mol, 5.92 ± 0.09 kcal/mol and 4.02 ± 0.10 kcal/mol for simulations with gHBfix_1-0_, gHBfix_0.5-0.5_, gHBfix_1-0__NBfix_0BPh-_pur__ and gHBfix_0.5-0.5__NBfix_0BPh-_pur__ potentials, respectively, comparable to those obtained from the initial 5 μs-long simulations (Table 2). This test, where two completely independent simulations performed using slightly different *ffs* produce very similar results upon reweighting, provides an additional confirmation that the 5 μs-long timescale is sufficient for the convergence of the ST-MetaD simulations for TLs (Figure 3). The NBfix_0BPh-_pur__ correction is either not affecting or slightly destabilizing the native state (Table 2).

**Table 2:**
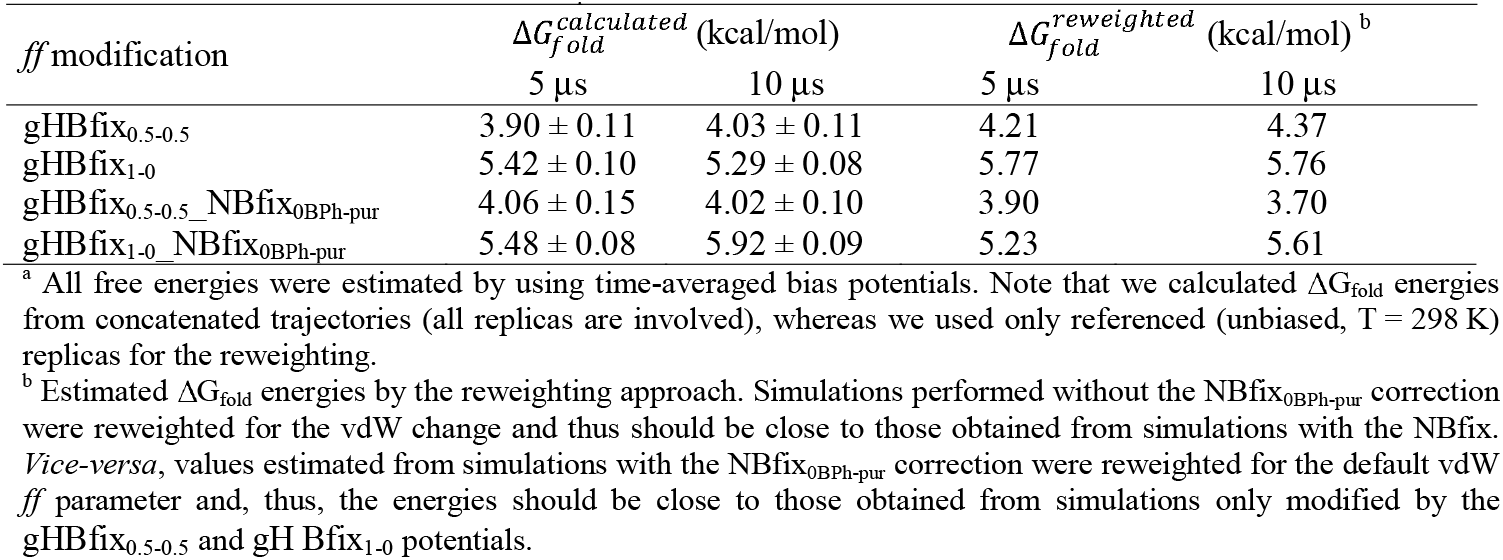
Comparison of directly calculated ΔG_fold_ values for r(gcUUCGgc) with values from the crossreweighting approach for the NBfixr_0BPh-_pur__ correction.^a^

The effect of the NBfix_0BPh-_pur__ correction for the r(gcUUCGgc) TL is complex. The correction is clearly supported by QM calculations, which identify excessive short-range repulsion of the –H8…O5’– interaction by the *ff*. The NBfix_0BPh-_pur__ correction stabilizes canonical A-form conformation for the G_S+1_ phosphate (the native conformation) and prevents flips to non-native *β_g+/γtrans_* G_S+1_ backbone conformation. However, despite the local stabilization of native sugar – phosphate backbone conformation, it does not lead to overall stabilization of the TL. We suggest the following explanation. Besides stabilizing the backbone in the native state, the correction is also stabilizing the G_L4_ residue in misfolded / unfolded states mostly sampling *anti* conformation of G_L4χ_ dihedral. This is a consequence of the fact that the NBfix_0BPh-_pur__ correction is not affecting *syn* residues where the –H8…O5’– distance is significantly larger, thus leading to a general stabilization of *anti* with respect to *syn*. Thus, the NBfix_0BPh-_pur__ correction accidentally stabilizes the state with G_L4_ bulging out ^21, 31, 32^ from its TL binding pocket. We can also explain the previous counter-intuitive observation that the NBfix_0BPh-pur_ correction speeds-up disruption of the UUCG TL native state in standard simulations initiated from the folded state. The key factor was not the G_L4_ 0BPh (as it is absent in the native state) but over-repulsive 0BPh interaction in a nearby G_S+1_ that slows down the disruption *kinetically;* for a detailed description see Ref. ^32^.

### Reweighting of additional *ff* modifications

The power of reweighting is to predict effects of *ff* changes without the necessity to perform additional extensive simulations, as demonstrated above for the NBfix_0BPh-_pur__ correction. Thus, we have used reweighting to test several simple *ff* modifications for both TLs. Namely, we investigated (i) destabilization of *syn* conformation of χ dihedral for adenines, (ii) stabilization of *syn* conformation of χdihedral for guanines, (iii) modification of vdW pair interactions for atoms participating in lone-pair…π contact between O4’ atom of ribose ring and atoms forming aromatic ring of nucleobases (sugar-base stacking), and (iv) modified 0BPh interaction of pyrimidine bases. The *ff* modifications were chosen based on preceding analyses of the UUCG TL^32, 96^ and also our earlier observation of higher (and spurious) propensity of adenines to sample *syn* states during tetranucleotide simulations.^77^ All results from reweighting are summarized in Table 3.

**Table 3:** Comparison of ΔG_fold_ energies obtained either from ST-MetaD simulations or from reweighting of respective simulations. ^a^

| System | r(gcGAGAgc) |  | r(gcUUCGgc) |  |  |  |  |  |
| --- | --- | --- | --- | --- | --- | --- | --- | --- |
| *ff* correction in simulations | gHBfix<sub>0.5-0.5</sub> | gHBfix<sub>1-0</sub> | gHBfix<sub>0.5-0.5</sub> | gHBfix<sub>1-0</sub> | gHBfix<sub>0.5-0.5</sub>—<br>NBfix<sub>0BPh-pur</sub> | gHBfix<sub>1-0</sub>—<br>NBfix<sub>0BPh-pur</sub> | gHBfix<sub>0.5-0.5</sub>—<br>NBfix<sub>0BPh-pyr</sub> | gHBfix<sub>1-0</sub>—<br>NBfix<sub>0BPh-pyr</sub> |
| $\Delta G_{\text{fold}}^{\text{calculated}}$ (kcal/mol) | 0.29 ± 0.17 | 0.14 ± 0.16 | 3.90 ± 0.11 | 5.42 ± 0.10 | 4.06 ± 0.15 | 5.48 ± 0.08 | 2.32 ± 0.11 | 4.67 ± 0.09 |
| *ff* parameter (reweighted by) <sup>b</sup> | $\Delta G_{\text{fold}}^{\text{reweighted}}$ (kcal/mol) | | | | | | | |
| **χ-dihedral** (*syn* minimum for A destabilized by 0.5 kcal/mol) | 0.24 | 0.12 | - | - | - | - | - | - |
| **0BPh_purines** ($R_{ij}$ for –H8...O5'– pair decreased by 0.25 Å) | -0.34 | -0.50 | 4.21 | 5.77 | - | - | 2.93 | 5.14 |
| **0BPh_pyrimidines** ($R_{ij}$ for –H6...O5'– pair decreased by 0.25 Å) | 0.13 | 0.05 | 3.06 | 4.37 | 3.65 | 4.78 | - | - |
| **0BPh_purines&pyrimidines** ($R_{ij}$ for both –H8...O5'– and –H6...O5'– pairs decreased by 0.25 Å) | -0.49 | -0.45 | 3.54 | 4.74 | - | - | - | - |
| **χ-dihedral** (*syn* minimum for G stabilized by 0.5 kcal/mol) | - | - | 3.92 | 5.32 | 4.00 | 5.31 | 2.10 | 4.51 |
| **lone-pair_π** ($R_{ij}$ for –O4'...C2– decreased by 0.25 Å) | - | - | 3.76 | 5.27 | 3.91 | 5.32 | 2.17 | 4.52 |
| **lone-pair_π** ($R_{ij}$ for –O4'...N1– decreased by 0.25 Å) | - | - | 3.87 | 5.38 | 4.01 | 5.43 | 2.26 | 4.62 |
| **lone-pair_π** ($R_{ij}$ for –O4'...N3– decreased by 0.25 Å) | - | - | 3.86 | 5.35 | 4.00 | 5.43 | 2.28 | 4.63 |

<sup>a</sup> Calculate $\Delta G_{\text{fold}}$ energies from respective ST-MetaD simulations are shown in the third row, whereas reweighted energies (for particular *ff* parameters, see the first column of the table) are displayed in the remaining part of the table. Energies were estimated from 5 μs-long simulations using time-averaged bias potentials. We calculated $\Delta G_{\text{fold}}$ energies from concatenated trajectories involving all twelve replicas, whereas we used only reference (unbiased) replicas for estimation of the reweighted energies. We note that calculated $\Delta G_{\text{fold}}$ values from the reference replica are very close (within 0.05 kcal/mol) from final values using all twelve replicas.
<sup>b</sup> Only vdW distances, i.e., $R_{ij}$ parameters, where the Lennard-Jones potential for interactions of atoms $i$ and $j$ is exactly zero, were modified. Depths of the potential well, i.e., $\epsilon_{ij}$ parameters, were kept at its default values for the respective atom pairs.

The (de)stabilization of *syn* state of the χ dihedral for purine bases seems to have no effect on the ΔG_fold_ data (Table 3). Also modification of pairwise vdW parameters allowing more compact lone-pair…π interaction characterizing the so-called Z-step motif^32, 96–98^ in UUCG TL did not affect reweighted ΔG_fold_ energies. The Z-step conformations contain sugar-base stacking with short distance between the O4’ and nucleobase rings for which QM calculations reveal that the *ff* overestimates the short-range repulsion of this interaction. We thus tested modified pairwise vdW parameters for –O4’…C2–, –O4’…N1–, and –O4’…N3– pairs and identified marginal ~0.2 kcal/mol stabilization of the native state only when adjusting the –O4’…C2– interaction (Table 3). It does not mean that we suggest sugar-base stacking is flawlessly described by the *ff* Rather, full inclusion of the QM vs. *ff* potential energy surface difference for the lone-pair…π interaction would require some more sophisticated modifications of the vdW term.^96^ MD description of Z-step conformations in DNA and RNA is a focus of ongoing research.^96, 99^

Reweighted ΔG_fold_ energies for modified 0BPh interactions, i.e., modified 0BPh for purine bases (NBfix_0BPh-_pur__), for pyrimidine bases (NBfix_0BPh-_pyr__), and simultaneous effect of both corrections, indicated stabilization of native states of both TLs. One exception is the standalone effect of the NBfix_0BPh-_pur__ correction for r(gcUUCGgc) TL, where we identified either comparable or slightly increased ΔG_fold_ energies by the NBfix. As discussed in the previous paragraph, it is caused by the rare G_L4_ *syn* conformation, where the NBfix_0BPh-_pur__ correction is rather stabilizing misfolded (bulged-out) ^21, 31, 32^ states instead of the native state with G_L4_ located in its binding pocket. We recently proposed^32^ that the NBfix_0BPh-_pur__ modification could be generally correct and the UUCG TL is just a specific system where its favorable free-energy effect is cancelled out by some other conformations on the landscape. In line with this, ΔG_fold_ energies from reweighted r(gcGAGAgc) ST-MetaD simulations are visibly lowered by the NBfix_0BPh-_pur__ correction (Table 3).

The NBfix_0BPh-_pyr__ correction appears slightly and significantly lower the ΔG_fold_ energy for the GAGA and UUCG TL, respectively (Table 3). We also reweighted simulations with simultaneous inclusion of both NBfix_0BPh-_pur__ and NBfix_0BPh-_pyr__ corrections and observed that ΔG_fold_ energies are lower by ~0.6 kcal/mol for both TLs (Table 3). As the reweighting might be affected by large statistical errors if the original and reweighted ensembles are not sufficiently overlapping (see Methods). We thus carried out r(gcUUCGgc) NBfix_0BPh-_pyr__ ST-MetaD simulations with gHBfix_1-0_ and gHBfix_0.5-0.5_ potentials. The directly calculated ΔG_fold_ energies are significantly lower in comparison with energies obtained from simulations without any NBfix correction (by ~1.6 kcal/mol and ~0.8 kcal/mol for simulations with gHBfix_0.5-0.5_ and gHBfix_1-0_ potentials, respectively, Table 1). Hence, the effect of NBfix_0BPh-_pyr__ correction on stability of the UUCG native state in full ST-MetaD runs is even higher than suggested from the reweighting (Table 3).

The above ST-MetaD data, nevertheless, show one surprising result. The NBfix_0BPh-_pyr__ correction improves the folding considerably more (difference of 0.8 kcal/mol) when being added to the gHBfix_0.5-0.5_ than to gHBfix_1.0_. This may indicate some kind of uncertainty in the ST-MetaD simulations. As the *ff* is supposed to be additive, effects from the gHBfix potential and NBfix_0BPh_ corrections to the ΔG_fold_ energy should be effectively summing up. The identified discrepancy is likely related to the way how the free energy landscape is explored in initial parts of the simulation, i.e., it could be side-effect of averaging of the bias potential. The low statistical errors of ST-MetaD simulations (Tables 1 and 2) can thus underestimate the uncertainty. Identification of true inaccuracies would, probably, require to carry out several independent ST-MetaD runs with same *ff* setting.^19, 20^ We plan to address this point is some future studies.

In summary, modification of 0BPh interactions appears to be a promising general RNA *ff* correction. It stabilizes native states of both most common RNA TLs and shifts the ΔG_fold_ values in the right direction. Further analysis of the effect of the NBfix_0BPh_ correction on other RNA systems is under way in our laboratories.

### 16×120 μs REST2 folding simulation of r(gcUUCGgc) TL shows no sign of convergence

In our earlier studies, ^24, 30, 74^ we often used the REST2 method, which worked excellently for the tetranucleotides. However, results from folding simulations of TLs were less clear.^30^ The simulations achieved some TL folding events, but analysis of overlaps of distributions between continuous (demultiplexed) replicas indicated a substantial lack of convergence, despite running quite extended REST2 runs. That is why in the present paper we replaced the REST2 protocol by the ST-MetaD one. As noted in the Introduction, efficiency of the RE methods is still a matter of debate.^14, 17–20, 64–68^ The sampling efficacy for a given system can be affected also by roughness of the free energy surface in a given *ff*.

In order to obtain more insights, we added one exceptionally long REST2 simulation of r(gcUUCGgc) TL (16 replicas with 120 μs per replica, i.e., more than 1.9 ms in total) into our portfolio of REST2 TL simulations.^30^ As simple gHBfix versions did not reveal any folding events during preceding REST2 simulations of UUCG TL (we even did not detect structures with successfully folded stem),^30^ we used a more complex version of the gHBfix potential prepared for the UUCG TL (gHBfix_UNCG19_, see Methods for details). In addition, we maintained setup from our earlier work^32^ and used the NBfix0BPh-pur correction with additional reduction of vdW radii of all non-polar H atoms (see Methods). This *ff* version provided promising results for folding of UUCG TL and is thus suitable for the test of the REST2 method.^32^

The primary goal of the long REST2 run was to understand the convergence behavior of the method for TLs. We detected only four complete folding events (i.e., from unfolded conformations to the native state), i.e., one event per ~0.5 ms of simulation time, each in one demultiplexed replica (Figure 4). The overall population of the entirely correct native state in the reference replica at 298 K was only ~5% while we obtained ~35% population of structures with the correct stems (Figures 4 and S4 in Supporting Information). This is in fact comparable with previously reported results from independent, significantly shorter (20 μs-long) simulation with exactly the same *ff* settings. However, neither significant time-prolongation nor slight increase in number of replicas (from 12^30^ to 16) did improve convergence as statistical errors remain very large (Table 1 and Figure S4 in Supporting Information). The result indicates that the sampling of the REST2 TL simulation is highly correlated and the effective sample size as characterized by completely independent and identically distributed conformations is insufficient.^19^ The prolonged REST2 does not show any sign of convergence improvement (Table S2 and Figures 4 and S4 in Supporting Information). Monitoring of the trajectories gives an impression that occurrence of a folded state in the ladder distorts and slows down exchanges across the ladder. Similar unsatisfactory convergence behavior can be inferred from the shorter GAGA REST2 simulations (see Figures S5 and S6 in Supporting Information).^30^

**Figure 4:**
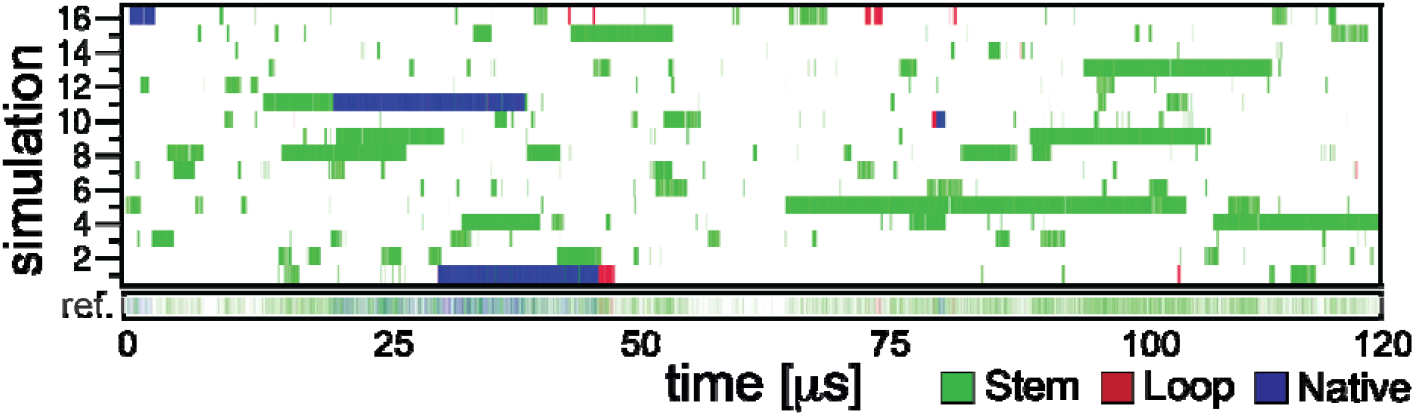
Conformational sampling of REST2 folding simulation of the r(gcUUCGgc) TL with the gHBfix_UNCG19_ correction in combination with the NBfix_0BPh_-pur__ correction and reduced vdW radii of all non-polar H atoms (see Methods). The panel shows time evolution of major conformers, i.e., (i) correctly folded A-form stem and loop (native states with all signature interactions formed, blue), (ii) folded A-form stem (loop not in native conformation, green), and (iii) correctly folded loop (stem not in A-form, red), in all sixteen continuous (demultiplexed) trajectories and the reference replica. Total population of the native state over the whole simulation in the reference replica is ~5%, which is still less than expected from experiments (~25%).^41–43^ Most importantly, however, dominant contribution to the population comes from the first 50 μs while the native state almost vanishes after this, illustrating the fundamental convergence problem.

### ST-MetaD dramatically outperforms REST2 in simulations of RNA TLs

Folding simulations of RNA TLs reported above were dominantly done by the combined ST-MetaD approach. Transitions between native and unfolded / misfolded states were facilitated by the MetaD using εRMSD^83^ from the native state as the CV. The REST2 protocol was applied to enhance the other degrees of freedom.

For the ST-MetaD, bias potentials are fluctuating in spite of the fact that the updated rate of the bias potential is decreasing over the time in well-tempered MetaD. Instantaneous ΔG_fold_ energies calculated from various simulation lengths thus fluctuate as well. Hence, we rather used time-averaged bias potentials, which were shown to speed-up the convergence significantly.^88^ It appears that ΔG_fold_ energies are converging quite quickly (Figures 2 and 3) and estimated statistical errors are ~0.2 kcal/mol on 5 μs time scales (Table 2). It is a major step forward compared to the sole REST2 simulations, which merely allowed to qualitatively assess if a given *ff* modification is or is not capable to spontaneously locate the native state when starting from the unfolded state.^30, 32^ As noted above, the true uncertainties of ST-MetaD simulations may be larger than suggested by the pure statistical errors. However, the ST-MetaD data appear to be quite robustly converged, concerning consistent results obtained from independent simulations performed with slightly different *ff*s.

An important factor to assess convergence of enhanced sampling simulations is monitoring of complete transitions between the folded and unfolded states, which are in detail summarized in the Tables S2 and S3 in Supporting Information. The ST-MetaD runs show dozens of such transitions which are typically by two orders of magnitude more frequent than in the REST2 simulations. The convergence as judged by monitoring the continuous (demultiplexed) replicas is not complete but quite reasonable, for full data see Figures S7-S15 in Supporting Information.

To achieve convergence in REST2 simulations of RNA TLs turns out to be a challenge. Although we have no comparison with standard simulations, the REST2 sampling enhancement appears to be low, if any. In our previous work, we used 10 μs-long REST2 simulations to assess how different gHBfix potentials affect folding of the r(gcGAGAgc) TL.^30^ We reported 20.0 ± 30.3 % and 9.1 ± 20.5 % populations of the native state in simulations with gHBfix_1-0_ and gHBfix_0.5-0.5_ potentials, respectively.^30^ The present ST-MetaD simulations give values of 44.13 ± 6.65 % and 37.97 ± 6.52 %, which are within the large statistical uncertainty of the REST2 data.

The huge REST2 statistical errors (note that the ranges formally include 0% population) obtained from the sophisticated bootstrapping protocol (see Supporting Information from Ref. ^30^ for details) clearly indicate that REST2 results for GAGA TL were not converged.^30^ It reflects the fact that we generated very few folding events to the native state, indicating severe lack of decorrelation in the REST2 runs. Complete folding – unfolding – folding transitions, which would be required to approach convergence,^19, 20^ were never detected in single 10 μs-long continuous replicas (Table S2 in Supporting Information). When we inspected time courses of the available REST2 simulations it appeared to us that the efficiency deteriorates when some folded molecule occurs in the ladder, as it tends to get stuck in the lower part of the ladder. Visits of the folded molecule in the upper part of the ladder are infrequent and very short, which may impair unfolding. Figures S16 and S17 in Supporting Information show climbing of continuous (demultiplexed) trajectories through the temperature ladder for r(gcGAGAgc) REST2 simulations.

Since the previous REST2 simulations were performed with 12 replicas,^30^ we carried out an additional REST2 simulation of r(gcGAGAgc) TL with gHBfix_1.0_ potential using 16 replicas (13 μs time per replica, Table S1 in Supporting Information). We obtained 19.20 ± 17.32 % population of the native state with only four folding events (Figure S18 in Supporting Information), confirming that slight increase in number of replicas is not providing any decisive convergence improvement for the REST2 method. The 16×120 μs r(gcUUCGgc) gHBfix_UNCG19_ REST2 simulation also reveals that neither adding few replicas nor prolonging the simulation time by an order of magnitude improves the convergence. One could, perhaps, add significantly more replicas in the ladder, but that would wipe out benefits of the REST2 method in comparison with other approaches, such as T-REMD or multidimensional replica exchange MD (M-REMD).^23^

Nevertheless, the obtained REST2 4.97 ± 5.65 % population of the native state would be so far the best result obtained for the UUCG TL 8-mer (when disregarding the huge statistical error again formally including 0% population). The observation of four folding events (each in a different continuous replica) to our opinion gives indication of some folding. However, the folded state is dominantly populated in the first 50 μs of the run. Thus, we performed also ST-MetaD simulation of r(gcUUCGgc) with the gHBfix_UNCG19_ potential together with the NBfix_0BPh-_pur__ and NBfix_0BPh-_pyr__ corrections (a comparable though not fully identical *ff* setting as used in the ultralong REST2 run, see Methods and Table 1). We obtained ST-MetaD ΔG_fold_ energy of 1.87 ± 0.12 kcal/mol (corresponding to 4.17 ± 0.82 % population of the native state, Table 1). It is actually in a good agreement with the REST2 data, though the agreement could be incidental.

The above result indicates that if the system is able to fold (having ΔGfold energy not far away from zero) the REST2 approach could provide at least some indication of ability of the system to fold. However, the primary quantity derived by REST2 simulation is the population of the folded state. Thus, the REST2 method cannot be used to estimate folding free energies of systems with too high or too low folding free energies, i.e., when population of one of the states (folded or un(mis)folded state) is negligible. With ΔGfold energy values above ~2.5 kcal/mol or below ~−2.5 kcal/mol one of the states would be marginally sampled in REST2 simulations irrespective of the convergence issue. In contrast the ST-MetaD simulations provide directly ΔG_fold_ values via the bias potential and thus they might be used generally regardless of the particular value of ΔG_fold_ free energy. This is another and significant advantage of the ST-MetaD approach compared to pure RE protocols, and it is indeed the main reason why MetaD was added in Ref.^25^.

The conformational space sampled by the ST-MetaD simulations can be affected by the choice of the CV(s).^8^ It appears that for the RNA TLs the εRMSD CV sufficiently well captures the slowest degree of freedom. Then the added REST2 protocol is robust enough to take care about the faster degrees of freedom. Figures S7-S15 in Supporting Information document conformational transitions in terms of εRMSD from the native state in all performed ST-MetaD simulations. We inspected conformations sampled during our ST-MetaD simulations and we found a significant amount of diverse misfolded states for both TLs. They include even the left-handed Z-form helix conformation^100^ (stem guanines in *syn* orientation, Figures S19 and S20 in Supporting Information). It indicates that the ST-MetaD approach is able quite broadly sample misfolded states and is not merely sampling the space between the native conformation and the unstructured part of the free energy landscape.

In summary, considering results presented here and in our earlier work,^30^ the ST-MetaD with εRMSD CV is strikingly more efficient compared to REST2 for RNA TLs. We note that this outcome may be system-specific, as performance of diverse enhanced-sampling methods is system dependent. Thus, for tetranucleotides, the REST2 method provides very satisfactory description,^30, 77^ while there is no guarantee that the ST-MetaD approach is a panacea for all other RNA systems.

## CONCLUDING REMARKS

Despite continuous development of RNA *ffs*, their performance is still problematic. Even small motifs like RNA TLs present challenge as there is an intricate balance between stability of the native state and other un(mis)folded states. ^24, 25, 27, 30^ Here, we employed ST-MetaD enhanced sampling method, i.e., combination of REST2 and well-tempered metadynamics aproaches, and calculated folding free energies (ΔG_fold_) of two most common r(gcGAGAgc) and r(gcUUCGgc) RNA TLs. We investigated effects of several *ff* adjustments on ΔG_fold_ energies and identify those, which are altering ΔG_fold_ energies in favor of the native states. We also compared performance of ST-MetaD and REST2 approaches.

Use of gHBfix potential leads to significantly increased stability (by 1.6 kcal/mol and 2.5 kcal/mol for GAGA and UUCG TLs, respectively) of the native state in comparison with the standard χ_OL3CP_ AMBER RNA *ff* Basic gHBfix versions like those mainly tested here, i.e., gHBfix_1-0_ (marked in the literature as gHBfix19) and gHBfix_0.5-0.5_,^32, 77^ are sufficient to correct the fold/un(mis)fold free energy disbalance for the GAGA TL. On contrary, the calculated ΔG_fold_energies for UUCG TL remained in the positive territory (corresponding to sporadic occurrence of the native state at 300 K). We also tested more complicated gHBfix version for UUCG TL (gHBfix_UNCG19_ potential, further coupled with NBfix corrections noted below) using both ST-MetaD and REST2 approaches and obtained promising populations of the native state ~5 % (Table 1), which is, however, still probably too low compared to experiments.^41–43^ Hence, further tunning of the gHBfix potential could be vital in order to stabilize UUCG native state in χ_OL3CP_AMBER RNA *ff*. Note that the gHBfix_UNCG19_ potential has not been tested for any other systems, in contrast to the gHBfix19.

We also show that NBfix adjustment of vdW parameters of the 0BPh interaction for both purine and pyrimidine nucleotides, i.e., NBfix_0BPh-_pur__ and NBfix_0BPh-_pyr__ corrections, shifts ΔG_fold_energies in the right direction. Initial results from reweighting showed that both NBfixes together are stabilizing native states of GAGA and UUCG TLs by ~0.6 kcal/mol. Subsequent ST-MetaD simulations revealed that the effect on ΔG_fold_ energies could be even larger (Table 3). Thus, application of both corrections simultaneously (NBfix_0BPh_) appears to be a justified RNA *ff* refinement although a broad testing using wide range of RNA motifs has to be performed in order to exclude side-effects.

Finally, we demonstrate much better convergence of the ST-MetaD approach compared to the sole REST2 method. REST2 runs are typically characterized by very few folding events and those can increase correlations across the replica ladder. The folded molecules tend to get stuck in the lower part of the ladder with infrequent visits of the upper part of the ladder. On contrary, combined ST-MetaD approach shows more frequent (typically by two orders of magnitude) transitions between the unfolded and folded states. ST-MetaD appears to be ideal for testing effects of simple *ff* adjustments and sequence dependencies of TLs and comparable motifs on calculated ΔG_fold_ energies.

## Supporting information

Supporting Information of the article

## ASSOCIATED CONTENT

### Supporting Information

Supplementary data are available free of charge via the Internet at http://pubs.acs.org/ and containing supporting tables and figures (PDF).

## ACKNOWLEDGMENT

We would like to dedicate the paper to the memory of Professor Neocles B. Leontis, for his seminal contribution to RNA bioinformatics. J.S. thanks Neocles for many years of inspiring collaboration. This work was supported by the project SYMBIT reg. number CZ.02.1.01/0.0/0.0/15_003/0000477 financed by the ERDF (J.S.) and by Czech Science Foundation (20-16554S to V.M., J.S., 18-25349S to M.J., P.K., and P.B.).

