## Supporting Information of the article for "Towards Convergence in Folding Simulations of RNA Tetraloops: Comparison of Enhanced Sampling Techniques and Effects of Force Field Corrections"

**Table of Contents**

Supporting Tables ........................................................ - 2 -

Supporting Figures ........................................................ - 5 -

References ........................................................ - 26 -

### Supporting Tables

**Table S1:** Overview of all newly-performed enhanced sampling simulations of RNA TLs.[a]

| Method | TL | gHBfix [b] | other *ff* modification [c] | Initial time (μs) | prolongation to (μs) | # of replicas |
|---|---|---|---|---|---|---|
| ST-MetaD | r(gcGAGAgc) | $gHBfix_{0.5\text{-}0.5}$ | - | 5 | - | 12 |
| ST-MetaD | r(gcGAGAgc) | $gHBfix_{1\text{-}0}$ | - | 5 | - | 12 |
| ST-MetaD | r(gcUUCGgc) | $gHBfix_{0.5\text{-}0.5}$ | - | 5 | 10 | 12 |
| ST-MetaD | r(gcUUCGgc) | $gHBfix_{1\text{-}0}$ | - | 5 | 10 | 12 |
| ST-MetaD | r(gcUUCGgc) | $gHBfix_{0.5\text{-}0.5}$ | $NBfix_{0BPh\text{-}pur}$ | 5 | 10 | 12 |
| ST-MetaD | r(gcUUCGgc) | $gHBfix_{1\text{-}0}$ | $NBfix_{0BPh\text{-}pur}$ | 5 | 10 | 12 |
| ST-MetaD | r(gcUUCGgc) | $gHBfix_{0.5\text{-}0.5}$ | $NBfix_{0BPh\text{-}pyr}$ | 5 | - | 12 |
| ST-MetaD | r(gcUUCGgc) | $gHBfix_{1\text{-}0}$ | $NBfix_{0BPh\text{-}pyr}$ | 5 | - | 12 |
| ST-MetaD | r(gcUUCGgc) | $gHBfix_{UNCG19}$ [d] | $NBfix_{0BPh}$ [e] | 5 | - | 12 |
| REST2 | r(gcGAGAgc) | $gHBfix_{1\text{-}0}$ | - | 10 | 13 | 16 |
| REST2 | r(gcUUCGgc) | $gHBfix_{UNCG19}$ [d] | $NBfix_{0BPh\text{-}pur}$ [f] | 20 [g] | 120 | 16 |

[a] All simulations were run with the $\chi_{OL3CP}$ RNA *ff* with specific versions of gHBfix potential and (where relevant) modified pairwise vdW parameters (NBfix, see Methods in the Main text).

[b] Base – base interaction term was stabilized differently, i.e., either only –NH…N– interactions were stabilized by 1.0 kcal/mol ($gHBfix_{1\text{-}0}$, known also as gHBfix19 version) or both –NH…N– and –NH…O– interactions were stabilized by 0.5 kcal/mol ($gHBfix_{0.5\text{-}0.5}$). Additional weakening of sugar donor – phosphate acceptor H-bonds (i.e., –OH…bO/nbO– interactions) by 0.5 kcal/mol was included in both versions; see Methods in the main text for details.

[c] Nonbonded fix (NBfix) was applied to modify the pairwise vdW parameters. Namely, we reduced the minimum-energy distance of Lennard-Jones potential (i.e., $R_{i,j}$ parameter) for the –H8…O5'– and –H6…O5'– pairs (more precisely between the H5 – OR and H4 – OR atom types, see Methods in the main text).

[d] Complex version of the gHBfix potential suggested specifically for the UUCG TL folding, where we also strengthened the sugar donor – base acceptor H-bonds and weakened base donor – sugar acceptor and sugar – sugar H-bonds (see Methods in the main text and Ref. [1] for details).

[e] Both $NBfix_{0BPh\text{-}pur}$ and $NBfix_{0BPh\text{-}pyr}$ corrections were applied.

[f] In addition to the $NBfix_{0BPh\text{-}pur}$ correction, vdW radii of all non-polar H atoms were reduced (see Methods in the main text and Ref. [1] for details).

[g] Independent simulation (the same *ff* settings) to the one published in our previous work.[1]

**Table S2:** Number of folding and unfolding events in available REST2 simulations of GAGA and UUCG RNA TLs.[a]

| TL | gHBfix [b] | other *ff* modification [c] | # (fold → unfold) events [d] | # (unfold → fold) events [e] | # of rep. | Time per rep. (μs) | Events per time ($\mu s^{-1}$) [f] |
|---|---|---|---|---|---|---|---|
| r(gcGAGAgc) | $gHBfix_{0\_0}$ | - | 0 (0) | 0 (0) | 12 | 10 [A] | 0.000 (0.000) |
| r(gcGAGAgc) | $gHBfix_{0.5\_0}$ | - | 0 (0) | 1 (0) | 12 | 10 [A] | 0.008 (0.000) |
| r(gcGAGAgc) | $gHBfix_{1\_0}$ | - | 0 (0) | 1 (0) | 12 | 10 [A] | 0.008 (0.000) |
| r(gcGAGAgc) | $gHBfix_{0\_0.5}$ | - | 0 (0) | 0 (0) | 12 | 10 [A] | 0.000 (0.000) |
| r(gcGAGAgc) | $gHBfix_{0.5\_0.5}$ | - | 0 (0) | 1 (0) | 12 | 10 [A] | 0.008 (0.000) |
| r(gcGAGAgc) | $gHBfix_{1\_0.5}$ | - | 1 (0) | 5 (0) | 12 | 10 [A] | 0.050 (0.000) |
| r(gcGAGAgc) | $gHBfix_{0\_1}$ | - | 0 (0) | 0 (0) | 12 | 10 [A] | 0.000 (0.000) |
| r(gcGAGAgc) | $gHBfix_{0.5\_1}$ | - | 0 (0) | 3 (0) | 12 | 10 [A] | 0.025 (0.000) |
| r(gcGAGAgc) | $gHBfix_{1\_1}$ | - | 1 (0) | 1 (0) | 12 | 10 [A] | 0.017 (0.000) |
| r(gcGAGAgc) | $gHBfix_{1\_0}$ | - | 3 (0) | 4 (0) | 16 | 13 [B] | 0.034 (0.000) |
| r(gcGAGAgc) [g] | $gHBfix_{0\_0}$ | WRESP-EP charges | 19 (4) | 4 (0) | 16 | 20 [C] | 0.072 (0.013) |
| r(gcGAGAgc) | $gHBfix_{0\_0}$ | WRESP-EP charges | 2 (1) | 3 (1) | 16 | 20 [C] | 0.016 (0.006) |
| r(gcUUCGgc) | $gHBfix_{UNCG19}$ | - | 1 (0) | 1 (0) | 12 | 20 [D] | 0.008 (0.000) |
| r(gcUUCGgc) | $gHBfix_{UNCG19}$ | $NBfix_{0BPh\text{-}pur}$, non-polar-H scaled | 2 (0) | 2 (0) | 16 | 20 [D] | 0.013 (0.000) |
| r(gcUUCGgc) [g] | $gHBfix_{UNCG19}$ | $NBfix_{0BPh\text{-}pur}$, non-polar-H scaled, dihedral restraint | 17 (1) | 1 (0) | 16 | 20 [D] | 0.056 (0.003) |
| r(gcUUCGgc) | $gHBfix_{UNCG19}$ | $NBfix_{0BPh\text{-}pur}$, non-polar-H scaled | 4 (0) | 4 (0) | 16 | 120 [B] | 0.004 (0.000) |

[a] All simulations were run with the $\chi_{OL3CP}$ RNA *ff* in ~1 M KCl salt excess using the Joung-Cheatham ion parameters[2] and OPC water model[3] (see Methods in the main text).

[b] see Table S1 footnotes for details of the gHBfix functions.

[c] Modified charges, modification of pairwise vdW parameters by NBfix and dihedral restraints. Note that on top of the $NBfix_{0BPh\text{-}pur}$ correction, vdW radii of all non-polar H atoms were reduced (see Methods in the main text) in some simulations (non-polar-H-scaled note). In one UUCG TL simulation (dihedral restraint note), we tried to support the native $U_{L1}$(2'-OH)…$G_{L4}$(O6) H-bond by restraining the $U_{L1}$(C1'-C2'-O2'-HO2') dihedrals (see Ref. [1] for details).

[d] Estimated total number of complete unfolding events from the native state in all replicas, where both structured loop and short stem (two GC base pairs) were lost and single-stranded like state was populated. Values in parentheses indicate likely observation of independent 'fold → unfold → fold → unfold' scenario across continuous (demultiplexed) replicas.

[e] Estimated total number of complete folding events in all replicas from the unfolded single-stranded like state to the native structure. Values in parentheses indicate likely observations of independent 'unfold → fold → unfold → fold' scenario across continuous (demultiplexed) replicas.

[f] Estimated number of fold and unfold events (values in parentheses correspond to independent 'fold → unfold → fold → unfold' and 'unfold → fold → unfold → fold' scenarios, see [d] and [e]) per 1 μs, i.e., number of events divided by number of replicas and total simulations time (μs).

[g] All replicas initiated from the native state.

[A] Data taken from Kührová et al. JCTC 2019, 15, 5, 3288–3305.[4]

[B] New simulations performed for this work.

[C] Data taken from Janeček et al. JCTC 2021, 17, 6, 3495-3509.[5]

[D] Data taken from Mráziková et al., JCTC 2020, 16, 12, 7601-7617.[1]

**Table S3:** Number of folding and unfolding events during ST-MetaD simulations of GAGA and UUCG RNA TLs performed in this work.[a]

| TL | gHBfix [b] | other *ff* modification [c] | # (fold → unfold) events [d] | # (unfold → fold) events [e] | # of rep. | Time per rep. (μs) | Events per time ($\mu s^{-1}$) [f] |
|---|---|---|---|---|---|---|---|
| r(gcGAGAgc) | $gHBfix_{0.5\text{-}0.5}$ | - | 21 (11) | 32 (17) | 12 | 5 | 0.883 (0.467) |
| r(gcGAGAgc) | $gHBfix_{1\text{-}0}$ | - | 22 (10) | 26 (12) | 12 | 5 | 0.800 (0.367) |
| r(gcUUCGgc) | $gHBfix_{0.5\text{-}0.5}$ | - | 33 (22) | 39 (23) | 12 | 10 | 0.600 (0.375) |
| r(gcUUCGgc) | $gHBfix_{1\text{-}0}$ | - | 33 (23) | 40 (27) | 12 | 10 | 0.608 (0.417) |
| r(gcUUCGgc) | $gHBfix_{0.5\text{-}0.5}$ | $NBfix_{0BPh\text{-}pur}$ | 25 (15) | 30 (19) | 12 | 10 | 0.458 (0.283) |
| r(gcUUCGgc) | $gHBfix_{1\text{-}0}$ | $NBfix_{0BPh\text{-}pur}$ | 40 (26) | 41 (28) | 12 | 10 | 0.675 (0.450) |
| r(gcUUCGgc) | $gHBfix_{0.5\text{-}0.5}$ | $NBfix_{0BPh\text{-}pyr}$ | 25 (14) | 27 (15) | 12 | 5 | 0.867 (0.483) |
| r(gcUUCGgc) | $gHBfix_{1\text{-}0}$ | $NBfix_{0BPh\text{-}pyr}$ | 22 (12) | 24 (14) | 12 | 5 | 0.767 (0.433) |
| r(gcUUCGgc) | $gHBfix_{UNCG19}$ | $NBfix_{0BPh}$ | 23 (9) | 27 (13) | 12 | 5 | 0.833 (0.367) |

[a] All simulations were run with the $\chi_{OL3CP}$ RNA *ff* and ~1 M KCl salt excess using the Joung-Cheatham ion parameters[2] and OPC water model[3] (see Methods in the main text).

[b] see Table S1 footnotes for details of the gHBfix functions.

[c] Modification of pairwise vdW parameters by NBfix (for details see footnotes of Table S1).

[d] Estimated total number of complete unfolding events from native state in all replicas, where both structured loop and short stem (two GC base pairs) were lost and single-stranded like state was populated (i.e., states with εRMSD > ~1.8 from the reference). Values in parentheses indicate likely observation of independent 'fold → unfold → fold → unfold' scenario across continuous (demultiplexed) replicas.

[e] Estimated total number of complete folding events across all replicas from the unfolded single-stranded like state to the native structure (i.e., states with εRMSD < ~0.7 from the reference). Values in parentheses indicate likely observations of independent 'unfold → fold → unfold → fold' scenario across continuous (demultiplexed) replicas.

[f] Estimated number of both fold and unfold events (values in parentheses correspond to independent 'fold → unfold → fold → unfold' and 'unfold → fold → unfold → fold' scenarios) per 1 μs (for details see footnotes of Table S2).

### Supporting Figures

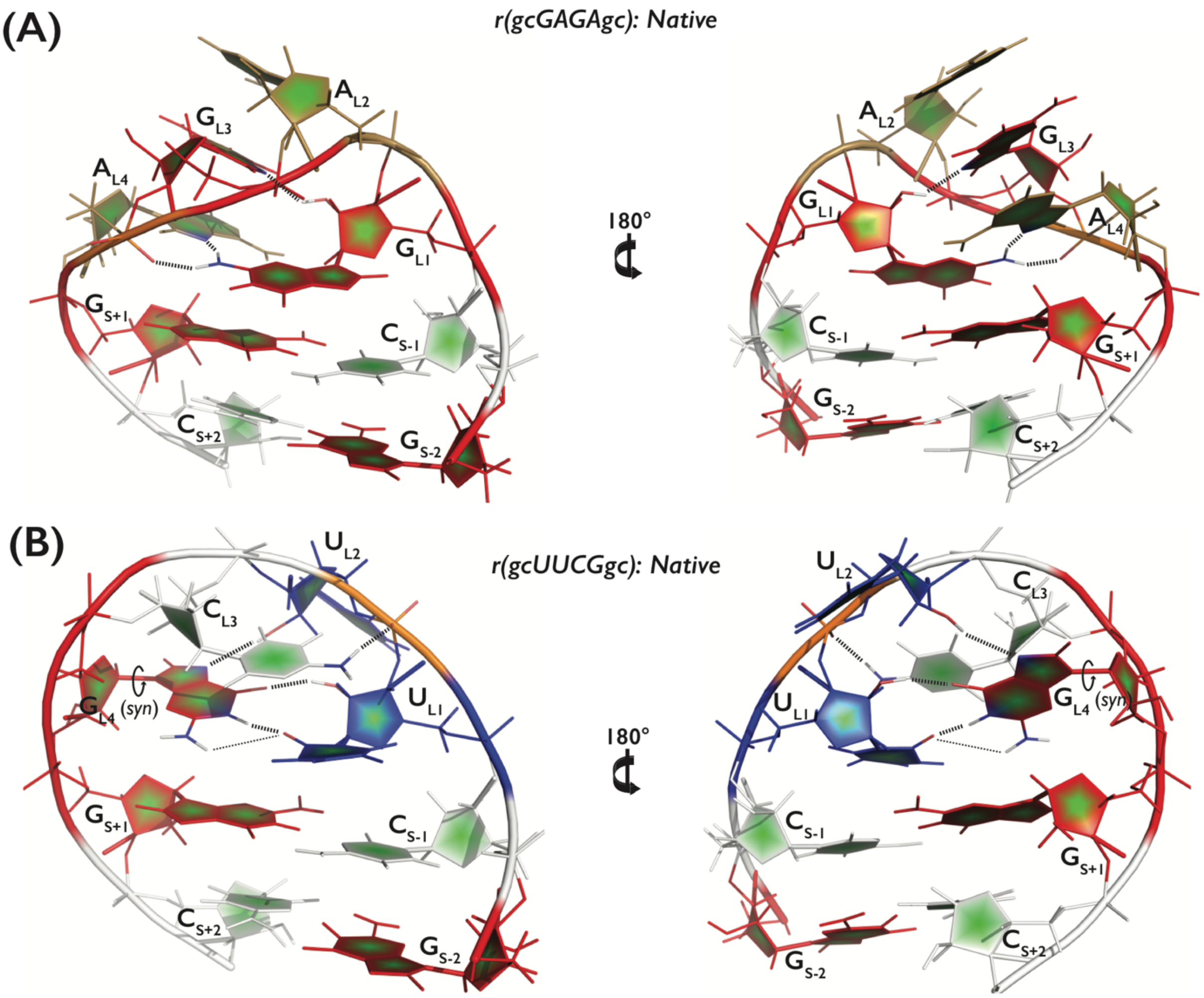


**Figure S1:** (A) r(gcGAGAgc) and (B) r(gcUUCGgc) RNA TLs in their native conformations. A, C, G and U nucleotides are colored in sand, white, red, and blue, respectively. Black dashed lines show the three and five signature (defining the consensus sequence) H-bonds in loops, i.e., (A) $G_{L1}$(N2H)…$A_{L4}$(*pro*-$R_P$), $G_{L1}$(N2H)…$A_{L4}$(N7), and $G_{L1}$(2'-OH)…$G_{L3}$(N7) for the GAGA TL and (B) $U_{L1}$(2'-OH)…$G_{L4}$(O6), $U_{L2}$(2'-OH)…$G_{L4}$(N7), $C_{L3}$(N4H)…$U_{L2}$(*pro*-$R_P$), and bifurcated $G_{L4}$(N1H/N2H)…$U_{L1}$(O2) for the UUCG TL, respectively.

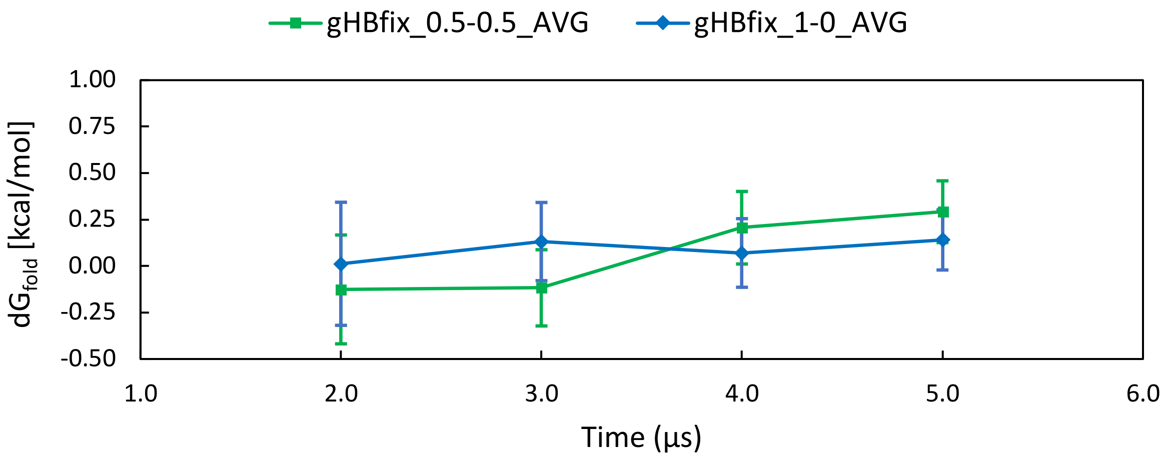


**Figure S2:** Comparison of $\Delta G_{fold}$ energies obtained from time-averaged bias potentials using data from 2 µs, 3 µs, 4 µs, and 5 µs-long ST-MetaD simulations of r(gcGAGAgc) TL. Statistical errors were obtained from concatenated trajectories and bootstrapping with 10, 12, 16, and 16 blocks for 2 µs, 3 µs, 4 µs, and 5 µs-long simulations, respectively. The plot compares behavior of two simulations modified by $gHBfix_{0.5\text{-}0.5}$ (green) and $gHBfix_{1\text{-}0}$ (blue) potentials.

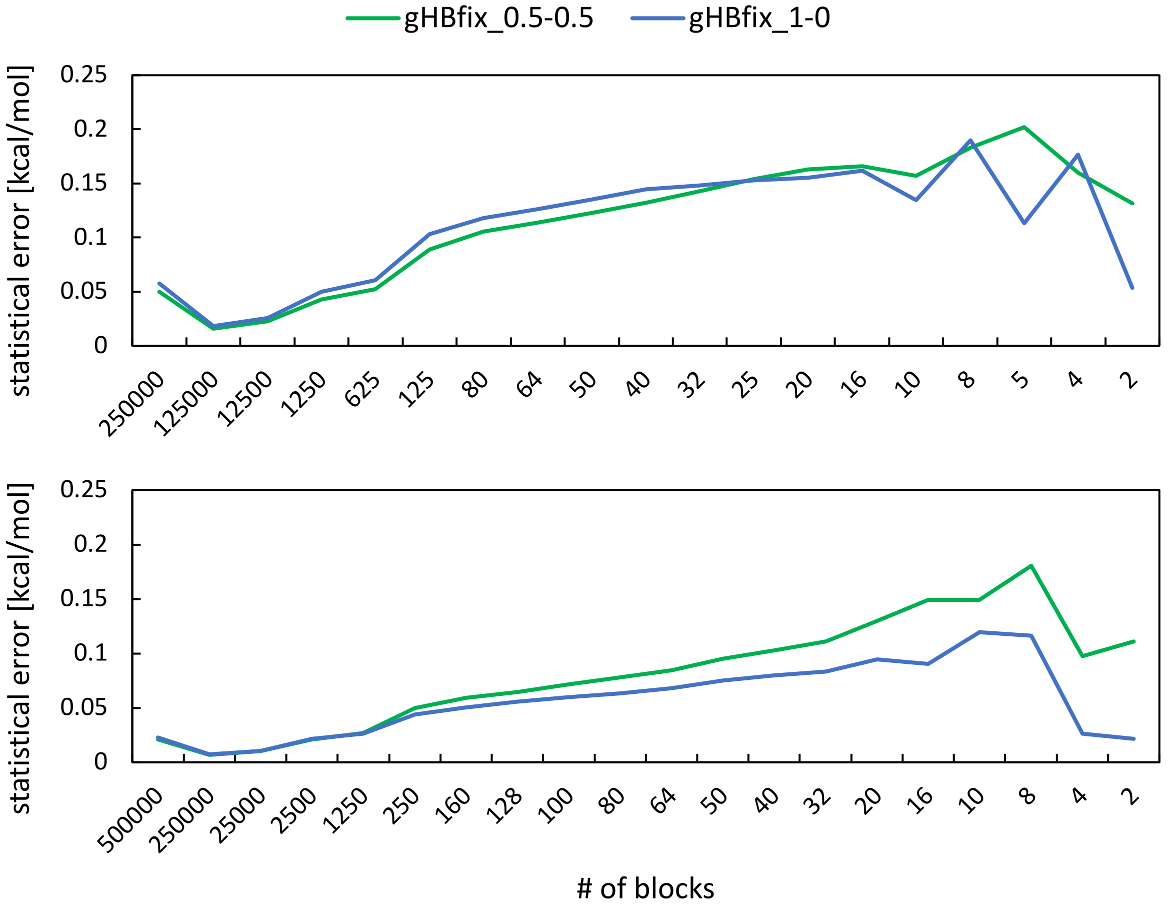


**Figure S3:** Fluctuations of statistical errors from bootstrapping and their dependence on the number of blocks. Plot on the top shows statistical errors from two r(gcGAGAgc) 5 µs-long ST-MetaD simulations with gHBfix$_{0.5\text{-}0.5}$ (green) and gHBfix$_{1\text{-}0}$ (blue) potentials, whereas bottom plot reveals errors from two r(gcUUCGgc) 10 µs-long ST-MetaD simulations again with gHBfix$_{0.5\text{-}0.5}$ (green) and gHBfix$_{1\text{-}0}$ (blue) potentials. Note that statistical errors for $\Delta G_{fold}$ energies reported in the main text used data from 16 and 32 blocks for 5 µs and 10 µs-long simulations, respectively.

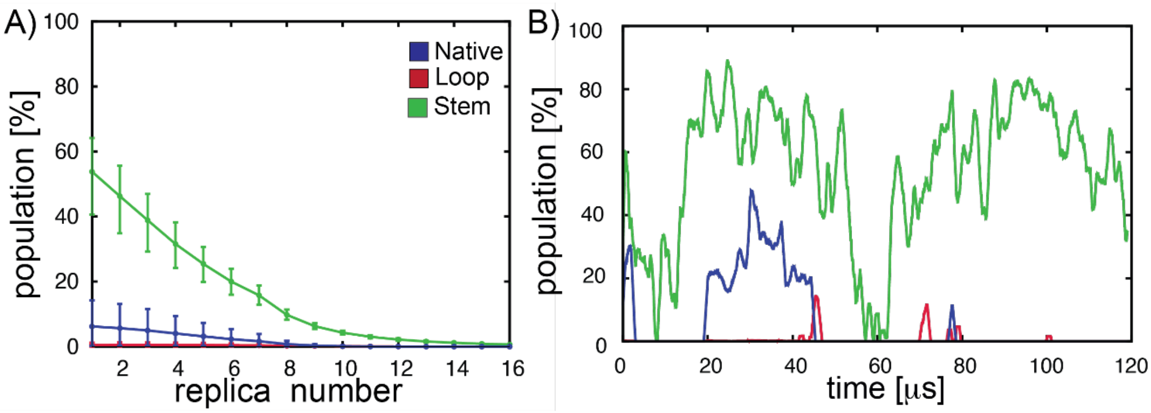


**Figure S4:** Convergence of the 120 µs-long REST2 folding simulation of the r(gcUUCGgc) TL with the gHBfix$_{UNCG19}$ correction in combination with the NBfix$_{0BPh\text{-}pur}$ correction and reduced vdW radii of all non-polar H atoms (Figure 4 in the main text). (A) Populations (%) of the most important types of structures, i.e., (i) correctly folded A-form stem and loop (native states with all signature interactions formed, blue), (ii) folded A-form stem (loop not in native conformation, green), and (iii) correctly folded loop (stem not in A-form, red), for all sixteen ladder replicas. Errors were estimated using sophisticated bootstrap implementation with resampling both over time- and replica-domains (see Ref. [4] for detailed description of the bootstrapping protocol). (B) Fluctuations of the most important types of conformers over the course of REST2 simulation obtained by time-averaging over the 1 μs window for the reference (unbiased, T = 298 K) replica (so that the value at 1 μs corresponds to the average over states populated between time of 0 to 1 μs).

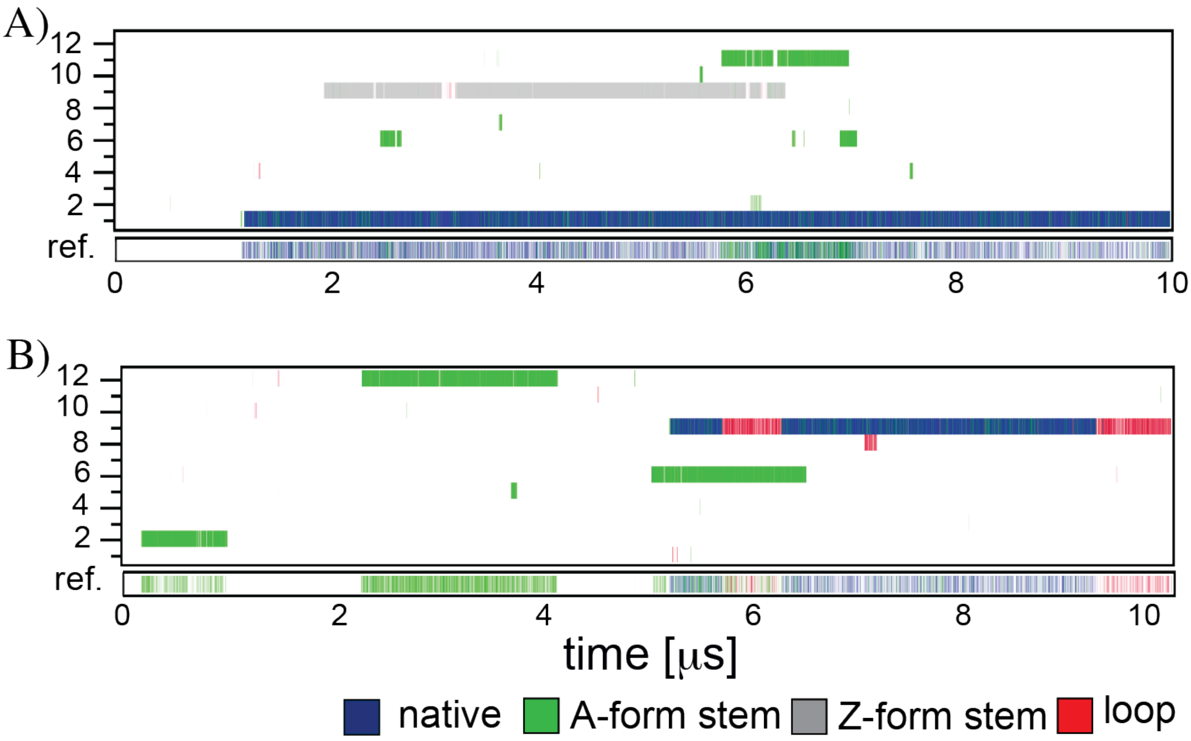


**Figure S5:** Conformational sampling of REST2 folding simulations of the r(gcGAGAgc) TL with the gHBfix$_{1\text{-}0}$ (A) and gHBfix$_{0.5\text{-}0.5}$ (B) potentials. Panel shows time evolution of major conformers in all twelve continuous (demultiplexed) trajectories and the reference replica (see Figure 4 in the main text). Both plots show just one complete folding event to the native state. Data were taken from Ref. [4].

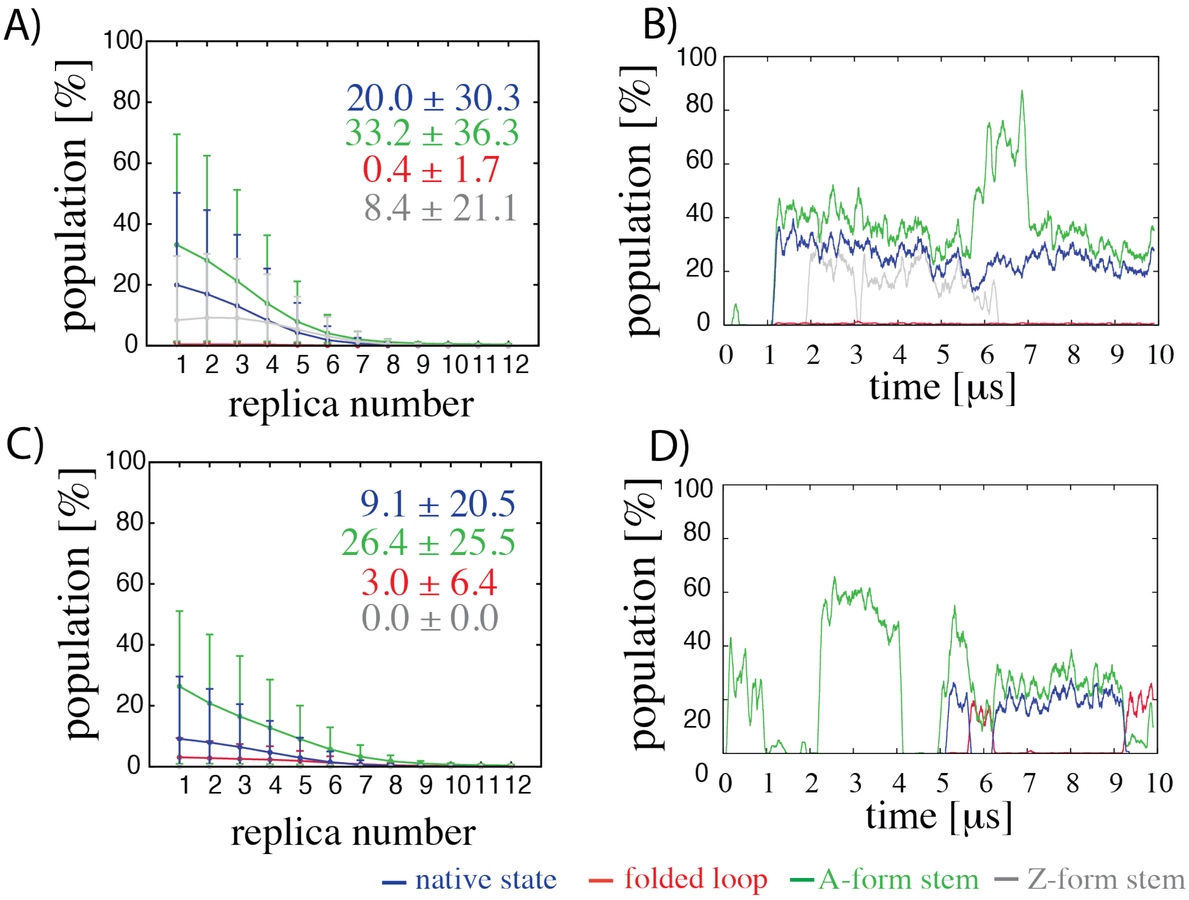


**Figure S6:** Convergence of REST2 folding simulations of the r(gcGAGAgc) TL with the gHBfix$_{1\text{-}0}$ (A, B) and gHBfix$_{0.5\text{-}0.5}$ (C, D) corrections (see Figure S5 for conformational sampling). (A) Populations (%) of the most important types of structures for all twelve ladder replicas (see Figure S4 for details). (B, D) Fluctuations of the most important types of conformers over the course of the REST2 simulations obtained by time-averaging over the 1 µs window for the reference (unbiased, T = 298 K) replica. Data were taken from Ref. [4].

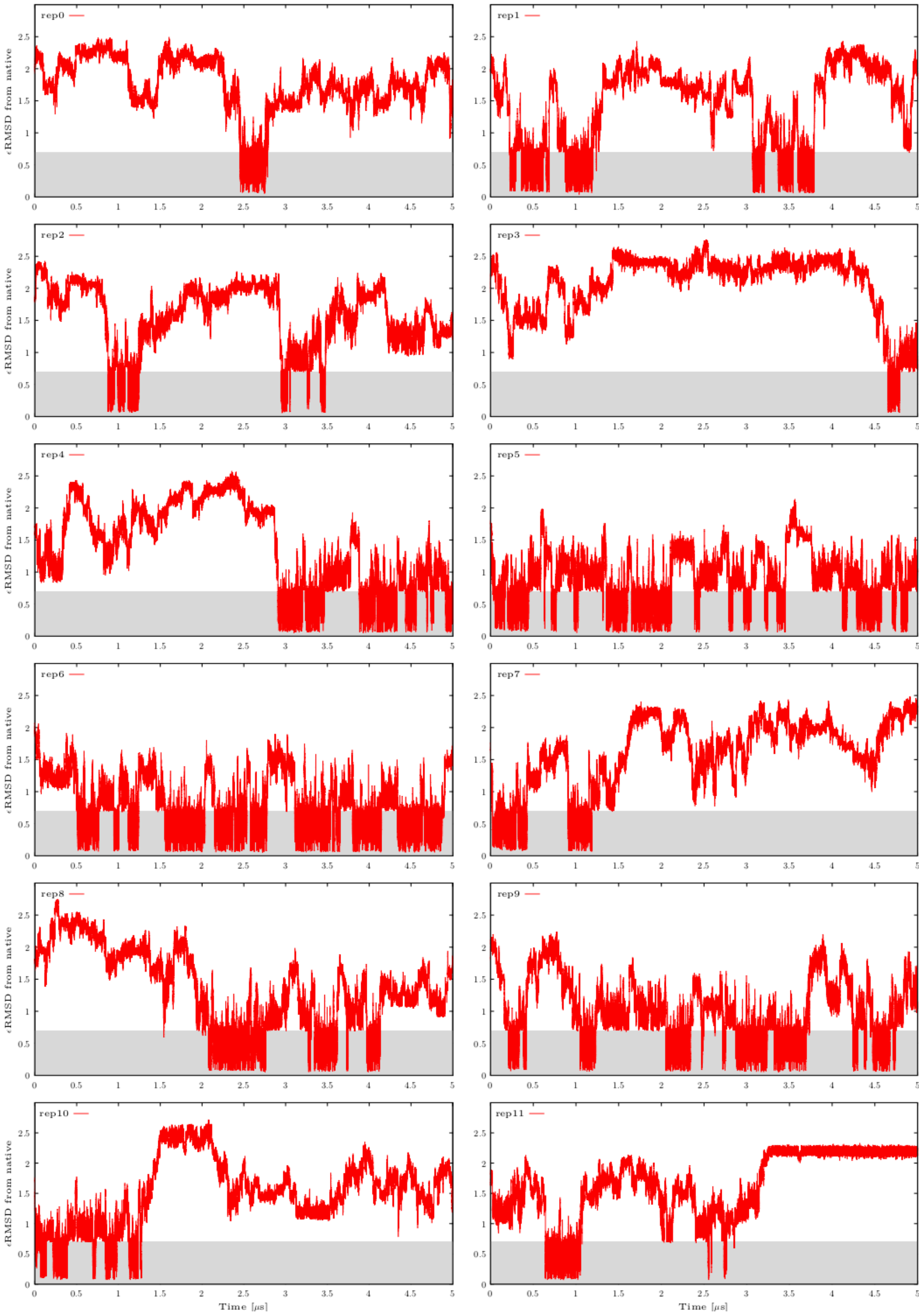


**Figure S7:** Calculated εRMSD from the native state in all twelve continuous (demultiplexed) replicas from ST-MetaD simulation of r(gcGAGAgc) TL with the gHBfix$_{1\text{-}0}$ potential. εRMSD values were calculated every 50 ps. The shaded area highlights states close to the reference (native state, εRMSD lower than 0.7).

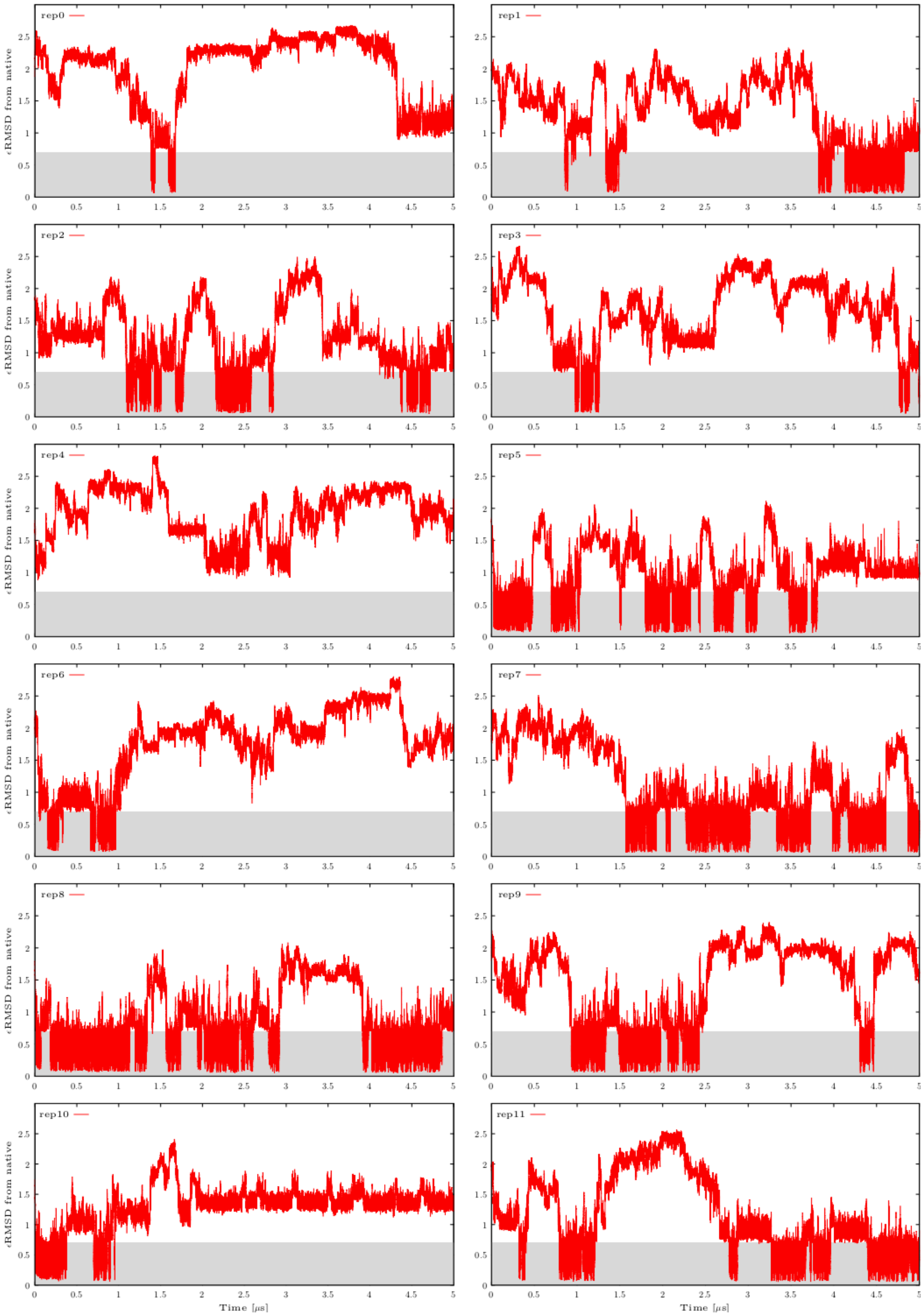


**Figure S8:** Calculated *ε*RMSD from the native state within all twelve continuous (demultiplexed) replicas from ST-MetaD simulation of r(gcGAGAgc) TL with the gHBfix$_{0.5\text{-}0.5}$ potential. See Figure S7 for more details.

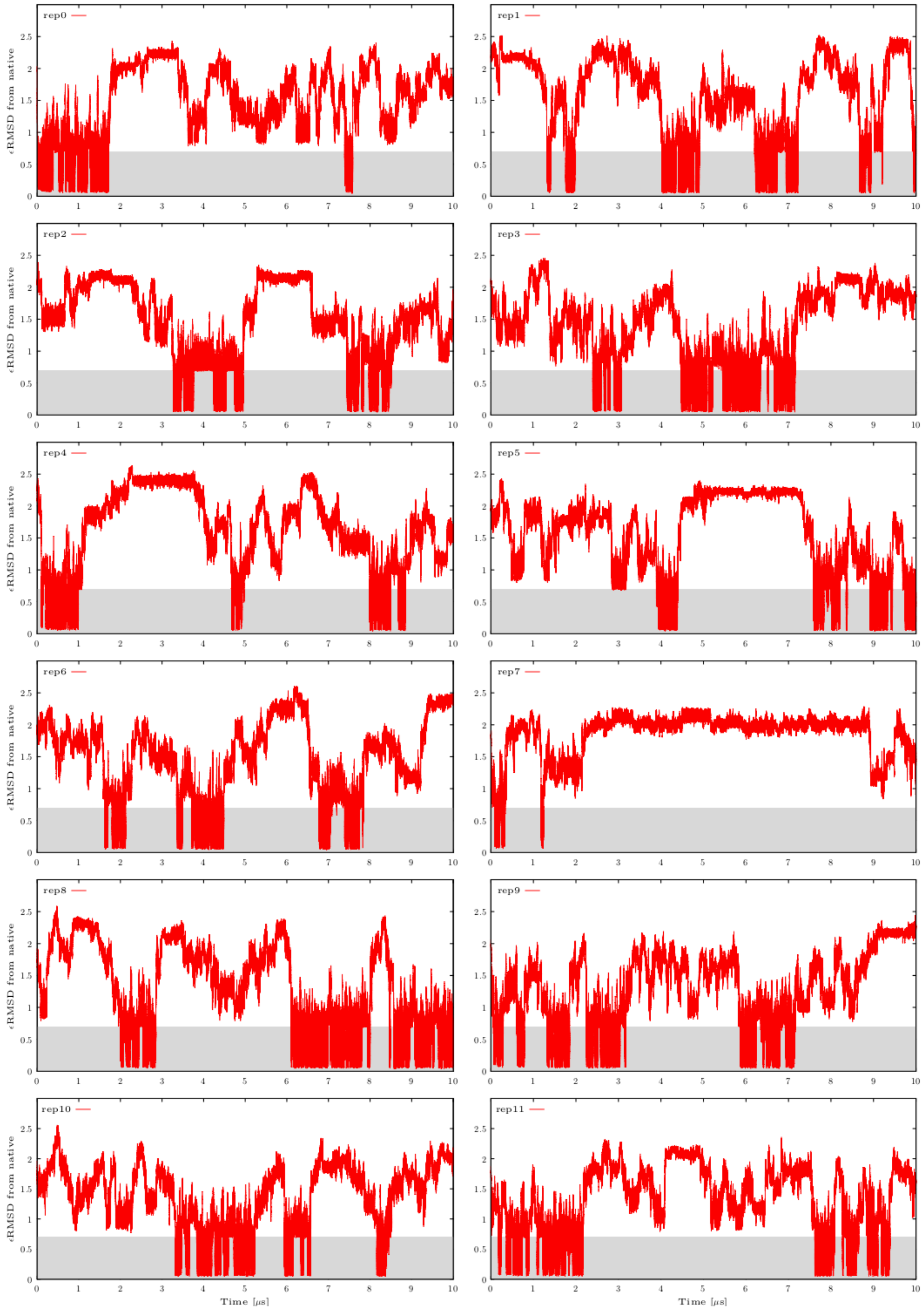


**Figure S9:** Calculated εRMSD from the native state in all twelve continuous (demultiplexed) replicas from ST-MetaD simulation of r(gcUUCGgc) TL with the gHBfix$_{1\text{-}0}$ potential. See Figure S7 for more details.

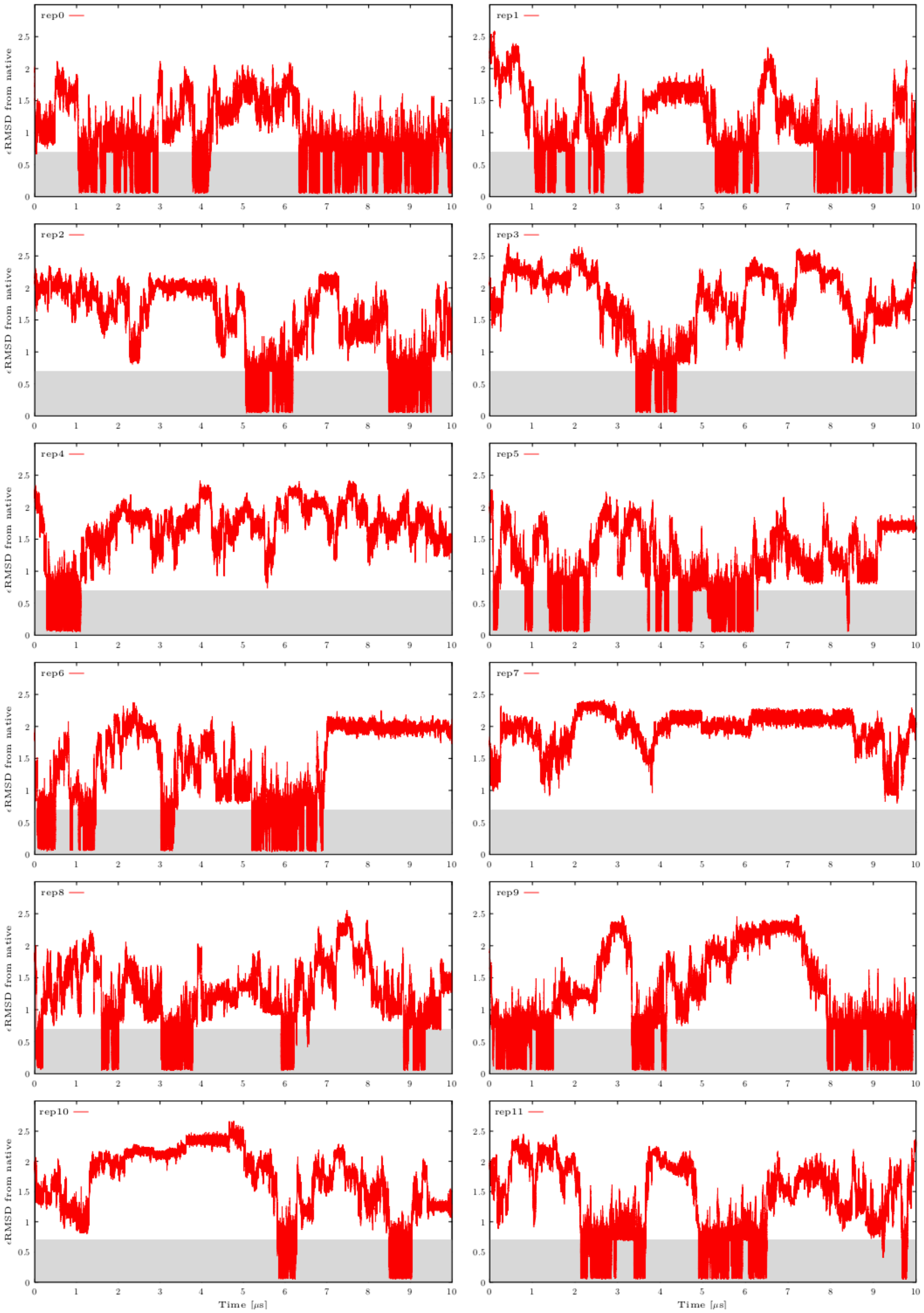


**Figure S10:** Calculated $\varepsilon$RMSD from the native state in all twelve continuous (demultiplexed) replicas from ST-MetaD simulation of r(gcUUCGgc) TL with the gHBfix$_{0.5\text{-}0.5}$ potential. See Figure S7 for more details.

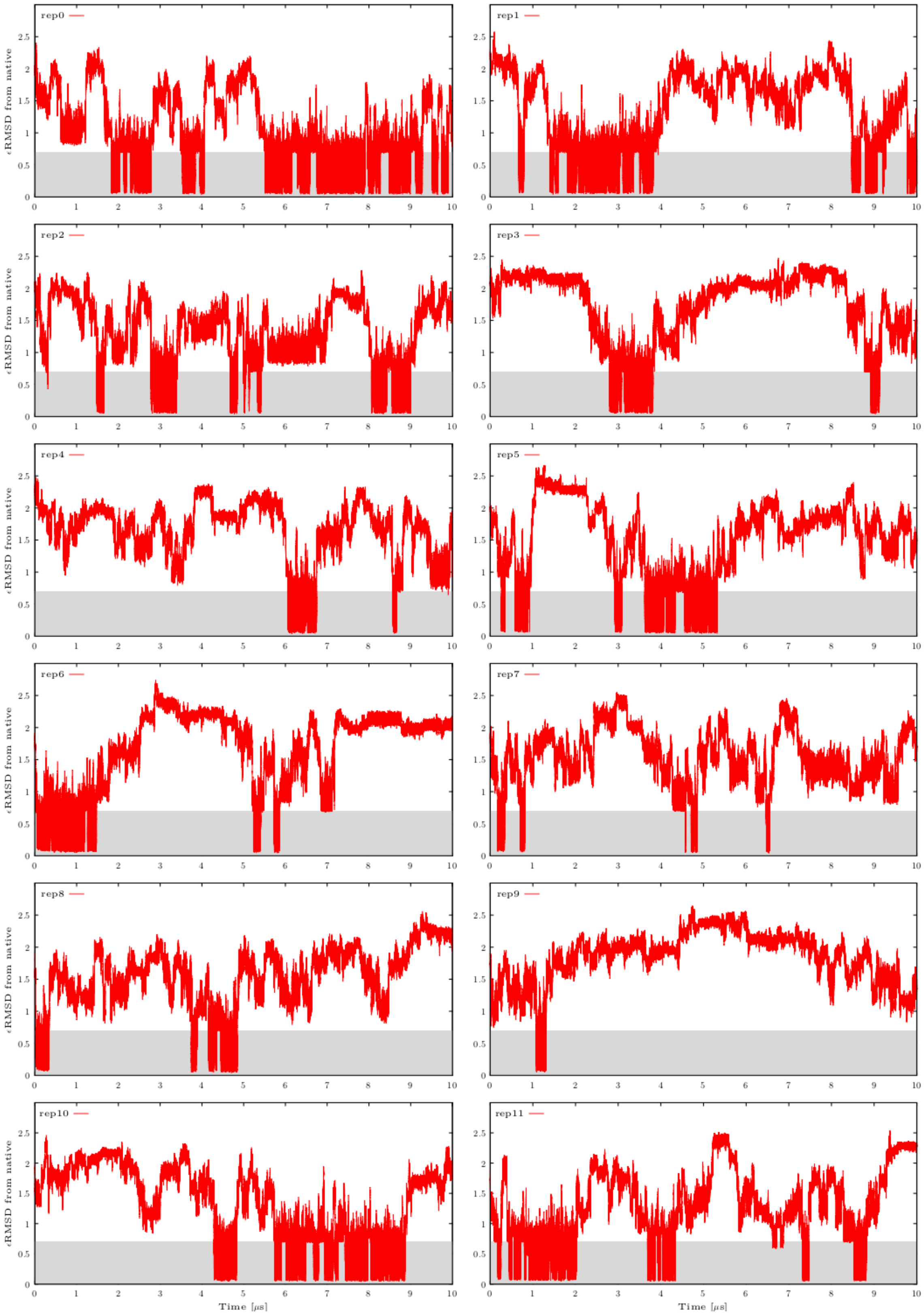


**Figure S11:** Calculated *ε*RMSD from the native state in all twelve continuous (demultiplexed) replicas from ST-MetaD simulation of r(gcUUCGgc) TL with the gHBfix$_{1\text{-}0}$_NBfix$_{0BPh\text{-}pur}$ potential. See Figure S7 for more details.

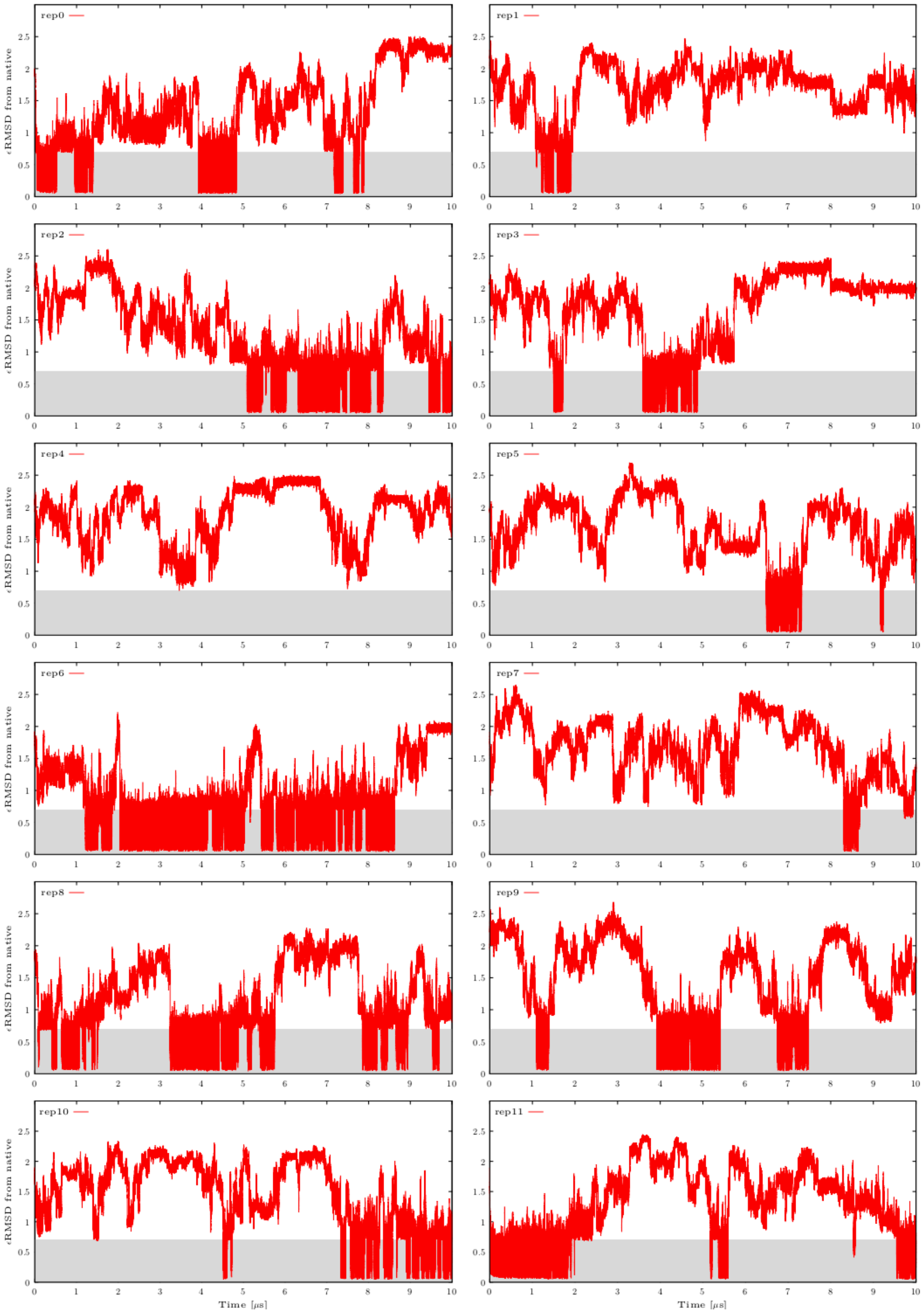


**Figure S12:** Calculated εRMSD from the native state in all twelve continuous (demultiplexed) replicas from ST-MetaD simulation of r(gcUUCGgc) TL with the gHBfix$_{0.5\text{-}0.5}$_NBfix$_{0BPh\text{-}pur}$ potential. See Figure S7 for more details.

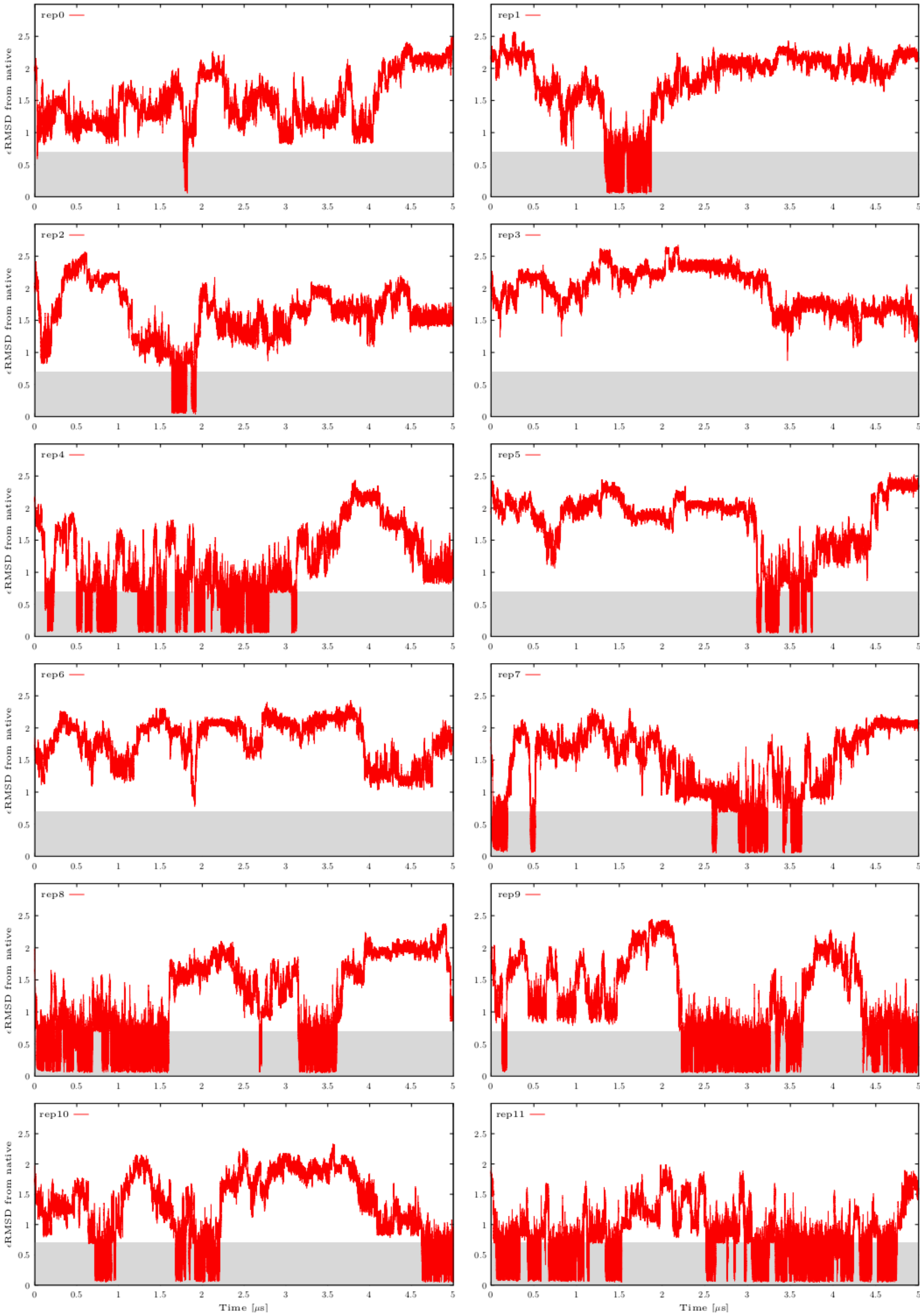


**Figure S13:** Calculated εRMSD from the native state in all twelve continuous (demultiplexed) replicas from ST-MetaD simulation of r(gcUUCGgc) TL with the gHBfix$_{1\text{-}0}$_NBfix$_{0BPh\text{-}pyr}$ potential. See Figure S7 for more details.

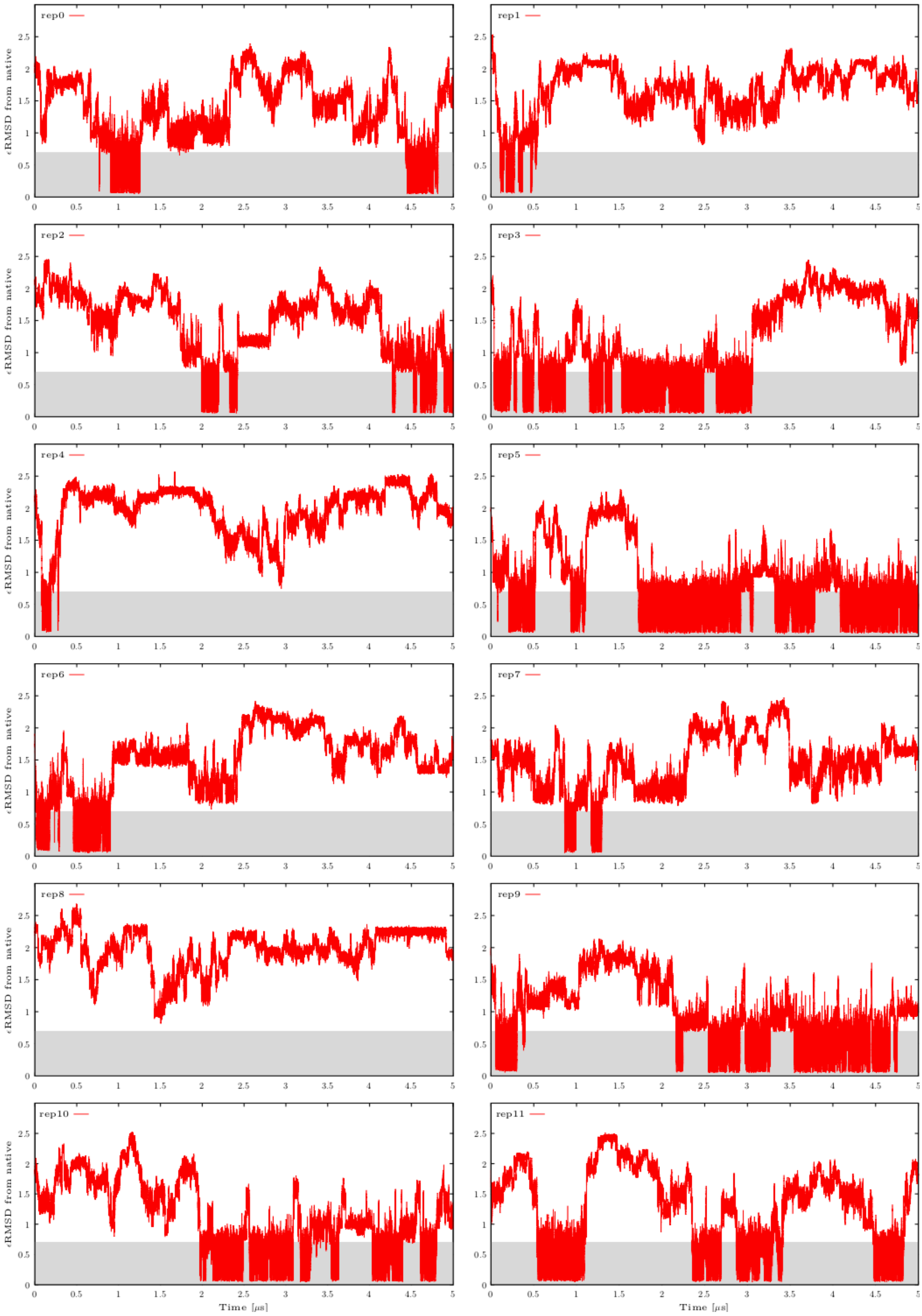


**Figure S14:** Calculated $\varepsilon$RMSD from the native state in all twelve continuous (demultiplexed) replicas from ST-MetaD simulation of r(gcUUCGgc) TL with the gHBfix$_{0.5\text{-}0.5}$_NBfix$_{0BPh\text{-}pyr}$ potential. See Figure S7 for more details.

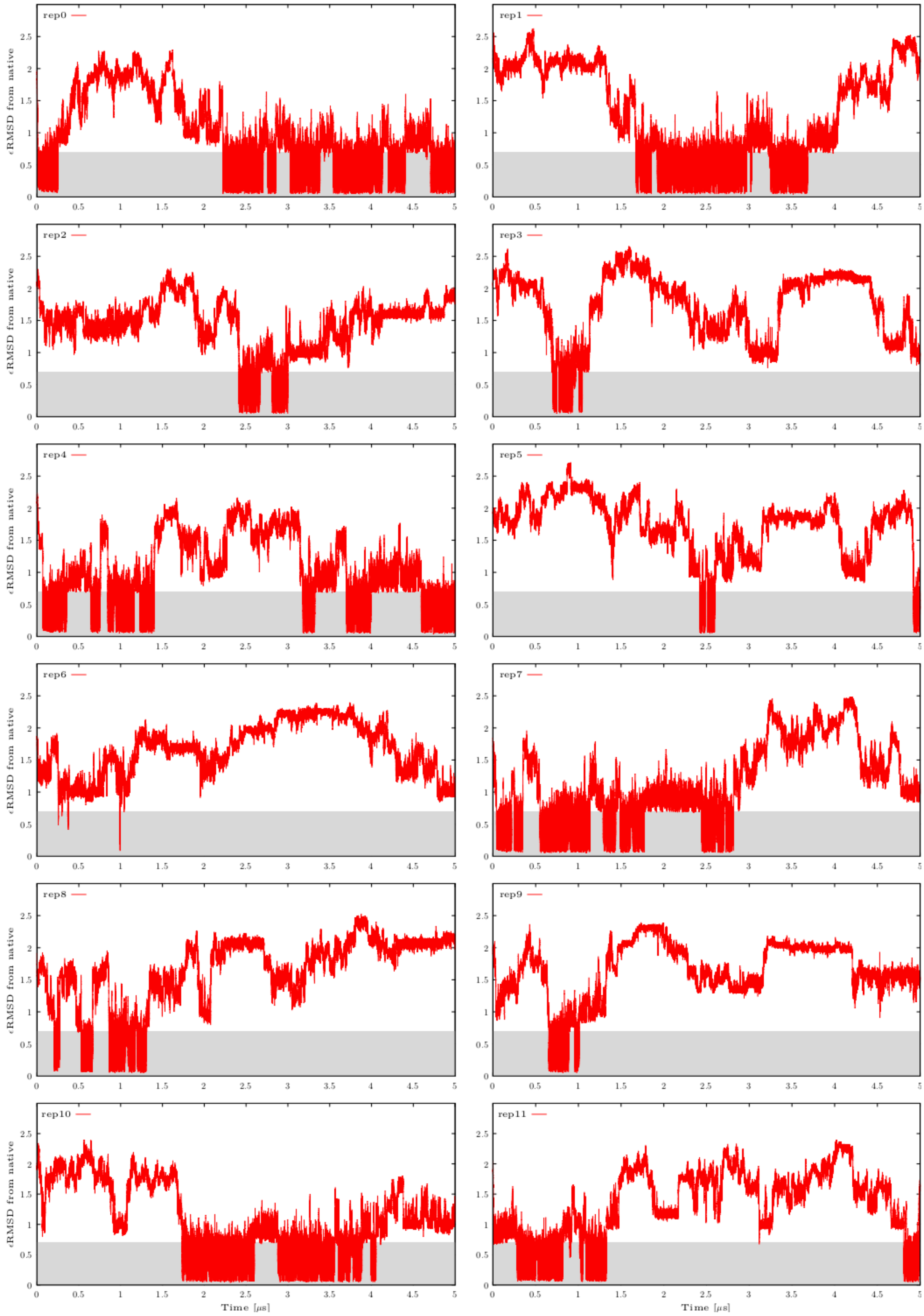


**Figure S15:** Calculated εRMSD from the native state in all twelve continuous (demultiplexed) replicas from ST-MetaD simulation of r(gcUUCGgc) TL with the gHBfix$_{UNCG19}$_NBfix$_{0BPh}$ potential. See Figure S7 for more details.

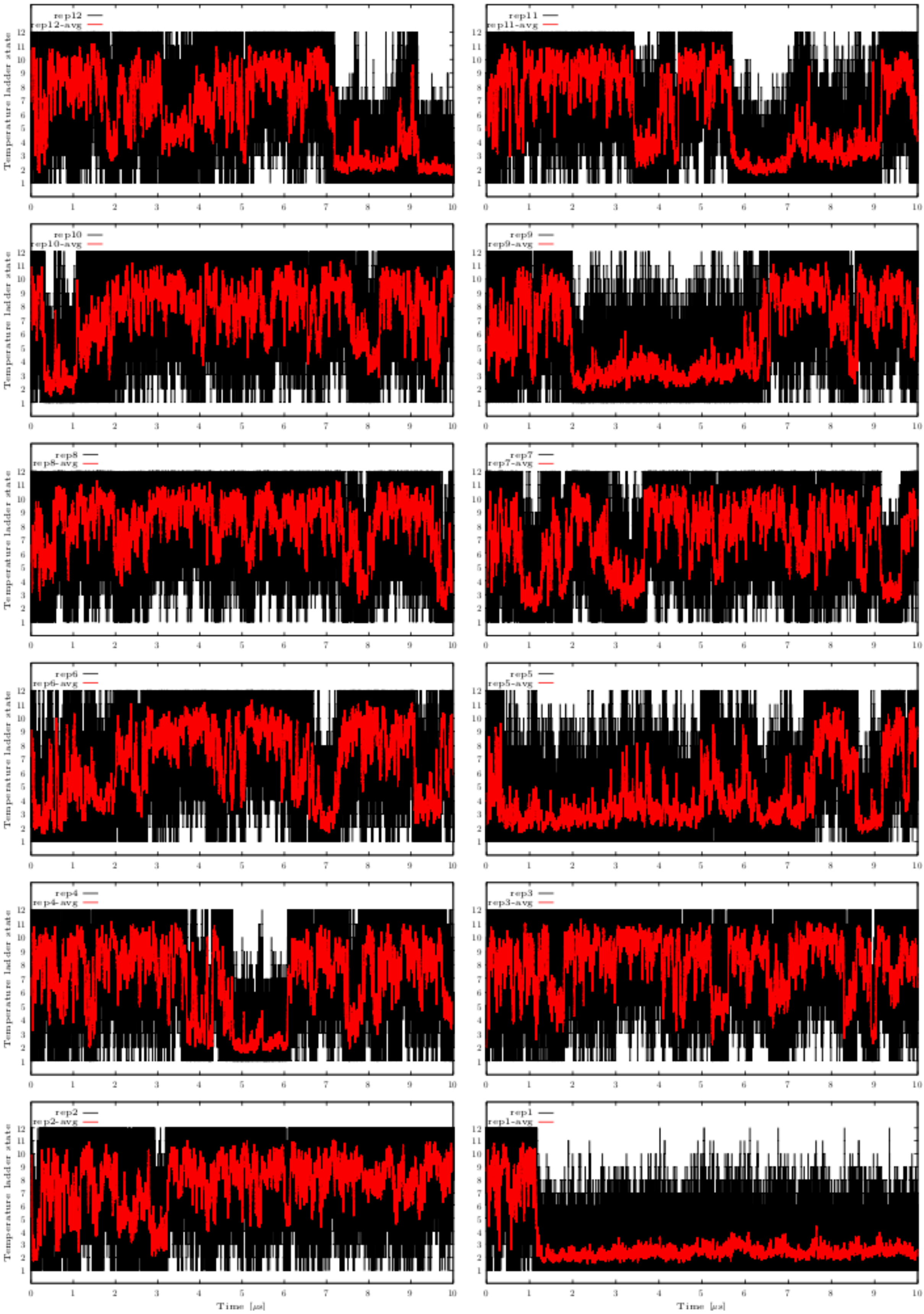


**Figure S16:** Movement of all twelve continuous (demultiplexed) trajectories through the temperature ladder from r(gcGAGAgc) TL REST2 folding simulation with the gHBfix$_{1-0}$ potential. Snapshots were saved every 10 ps (black lines) and smoothing (i.e., averaging over

100 consecutive snapshots, red lines) was also used for better visibility. Possible occurrence of folded states in the ladder can distort exchanges across the ladder, i.e., even though the replica with folded conformation is allowed to go up in the temperature ladder, it does no spend sufficient time there for successful unfolding. Simulation taken from Ref. [4]. Note that once the TL is folded in the continuous replica 1 (cf. also Figs S5 and S6), its capability to visit higher parts of the ladder is visibly suppressed.

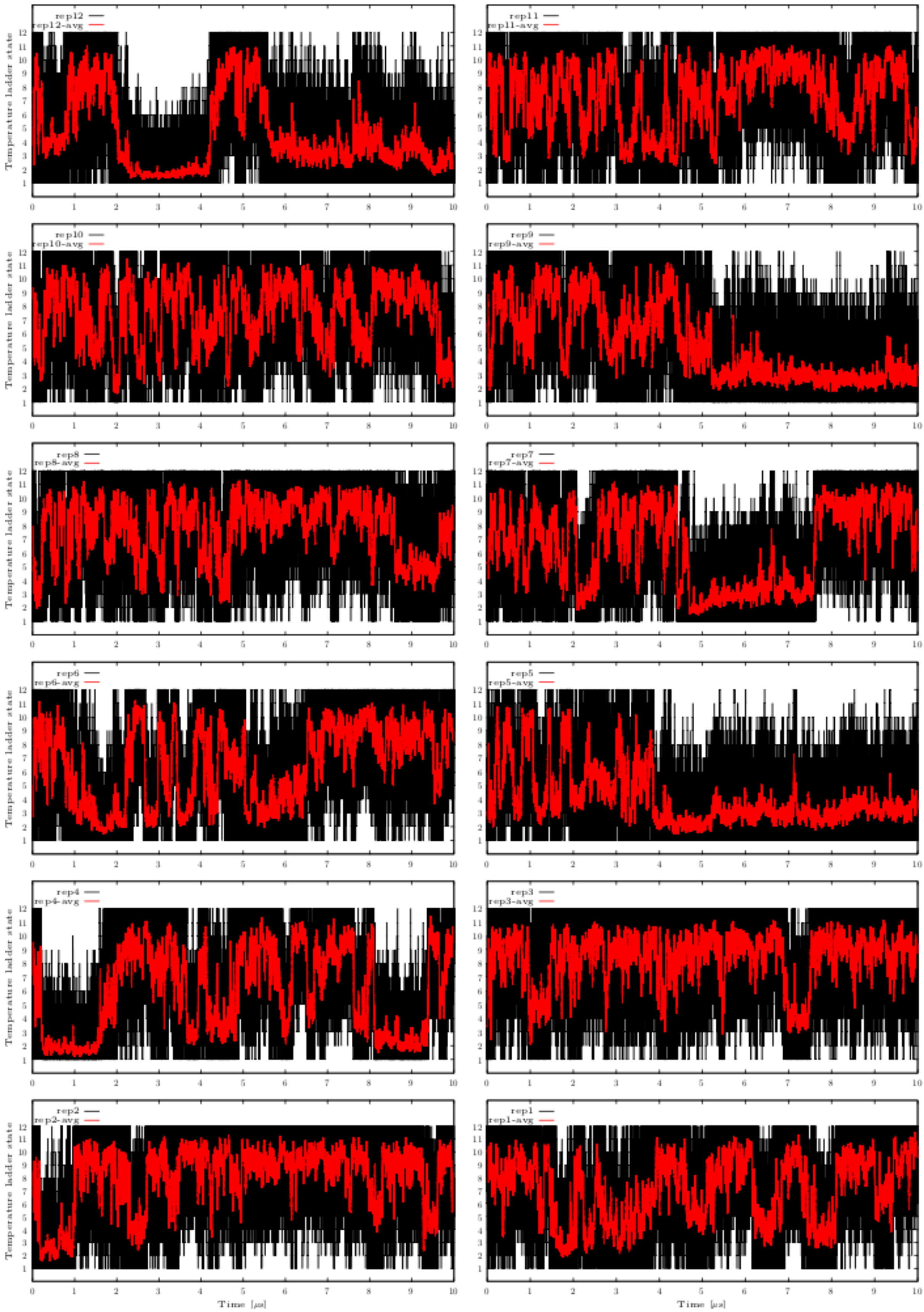

**Figure S17:** Movement of all twelve continuous (demultiplexed) trajectories through the temperature ladder from r(gcGAGAgc) TL REST2 folding simulation with the gHBfix$_{0.5\text{-}0.5}$ potential (see Figure S16 for details). Simulation taken from Ref. [4].

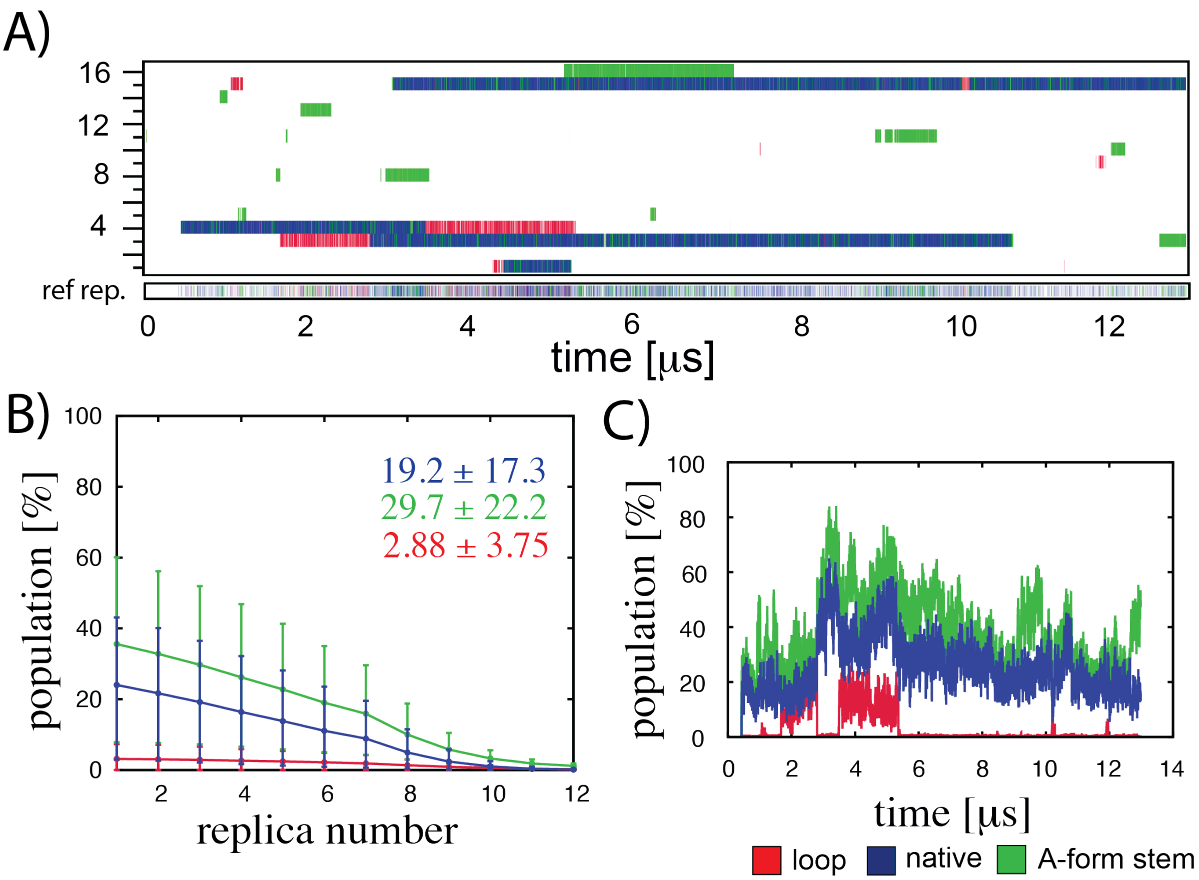


**Figure S18:** Conformational sampling of the 13 μs-long REST2 folding simulation of the r(gcGAGAgc) TL with the gHBfix$_{1\text{-}0}$ potential (16 replicas). (A) Panel shows time evolution of major conformers in all sixteen continuous (demultiplexed) trajectories and the reference replica. (B) Populations (%) of the most important types of structures (loop only, native state, stem only) for all sixteen ladder replicas. (C) Fluctuations of the most important types of conformers over the course of the REST2 simulation obtained by time-averaging over the 1 μs window for the reference (unbiased, T = 298 K) replica.

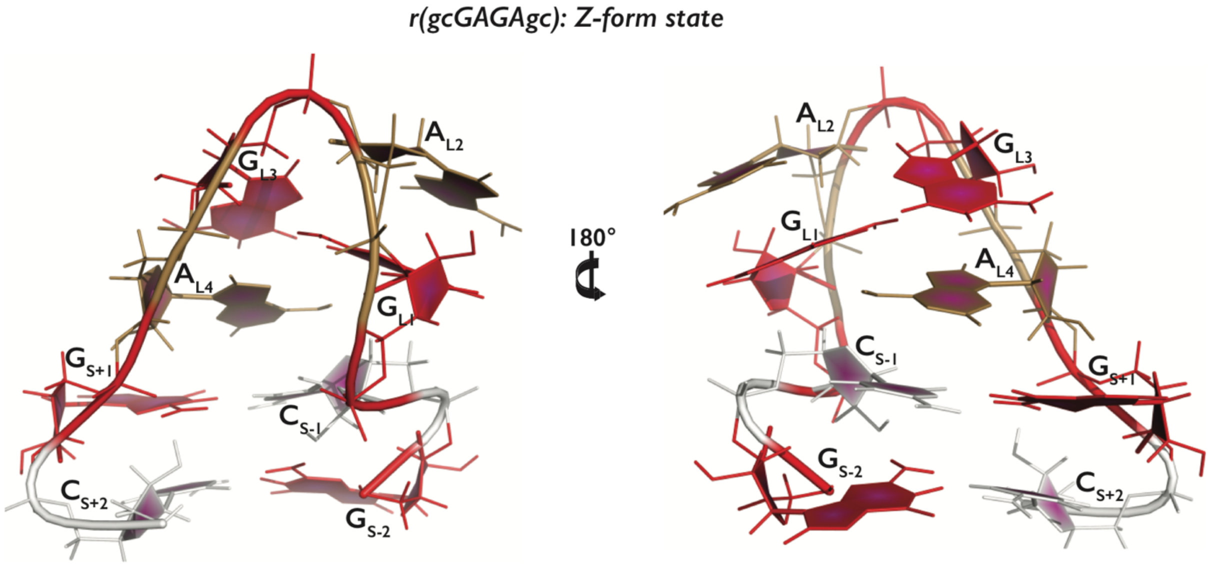


**Figure S19:** Tertiary structure of the r(gcGAGAgc) TL showing the left-handed Z-form helix conformation (G nucleotides in *syn* orientation). A, C, G and U nucleotides are colored in sand, white, red, and blue, respectively.

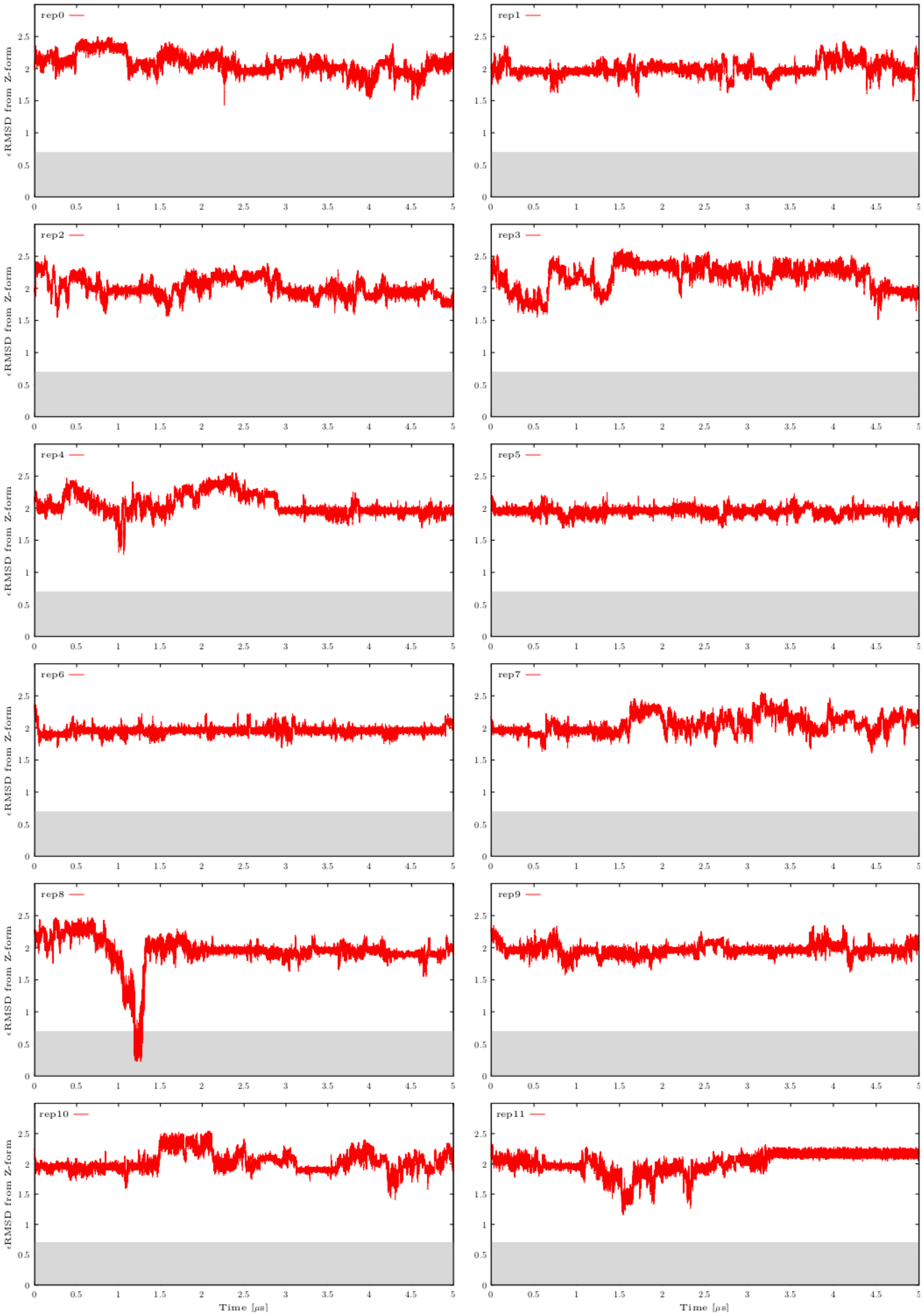


**Figure S20:** Calculated εRMSD from the Z-form state (see Figure S19 for the structure) in all twelve continuous (demultiplexed) replicas in one selected ST-MetaD simulation of r(gcGAGAgc) TL with the gHBfix$_{1\text{-}0}$ potential. εRMSD values were calculated every 50 ps.

The shaded area highlights states close to the reference (Z-form state, $\varepsilon$RMSD lower than 0.7). Plots show that the ST-MetaD approach is able to find states with Z-form but their formation is sporadic. This, however, is not a problem as this state is expected to be rare. We note that arrangement of nucleotides in the loop could be different that those shown at Figure S19 (from which the $\varepsilon$RMSD was calculated). However, we did not detect state with the native-like arrangement in the loop and Z-form stem.
